# Reassessing the validity of slow-wave dynamics as a proxy for NREM sleep homeostasis

**DOI:** 10.1101/748871

**Authors:** Jeffrey Hubbard, Thomas C. Gent, Marieke M. B. Hoekstra, Yann Emmenegger, Valerie Mongrain, Hans-Peter Landolt, Antoine R. Adamantidis, Paul Franken

**Affiliations:** Center for Integrative Genomics, University of Lausanne, Lausanne, Switzerland; Department of Neurology, Inselspital University Hospital, University of Bern, Bern, Switzerland; Department of Neuroscience, Université de Montréal, Montreal, QC, Canada; Institute of Pharmacology and Toxicology, University of Zürich, Zurich, Switzerland; Sleep & Health Zurich, University of Zürich, Zurich, Switzerland

**Keywords:** sleep homeostasis, delta power, active sleep, NREMS, physiology, fast/slow delta, cross-species

## Abstract

Sleep-wake driven changes in NREM sleep (NREMS) EEG delta (δ: ∼0.75-4.5Hz) power are widely used as proxy for a sleep homeostatic process. We noted frequency increases in δ-waves in sleep-deprived (SD) mice, prompting us to re-evaluate how slow-wave characteristics relate to prior sleep-wake history. We discovered two types of δ-waves; one responding to SD with high initial power and fast, discontinuous decay (δ2: ∼2.5-3.5Hz) and another unrelated to time-spent-awake with slow, linear decays (δ1: ∼0.75-1.75Hz). Human experiments confirmed this δ-band heterogeneity. Similar to SD, silencing of centromedial thalamus neurons boosted δ2-waves, specifically. δ2-dynamics paralleled that of temperature, muscle tone, heart-rate, and neuronal UP/DOWN state lengths, all reverting to characteristic NREMS levels within the first recovery hour. Thus, prolonged waking seems to necessitate a physiological recalibration before typical NREMS can be reinstated. These short-lasting δ2-dynamics challenge accepted models of sleep regulation and function based on the merged δ-band as sleep-need proxy.

## Introduction

Research on the function of sleep has been established on the premise that this behavioral state remedies the wear and tear caused by preceding waking. With the aim to gain insight into the underlying neuronal processes and anatomical pathways, researchers have focused on variables that accumulate during waking and dissipate across sleep, which could serve as proxy measures for what is referred to as sleep homeostasis. Nap and sleep deprivation (SD) studies in birds and mammals have demonstrated that EEG delta (∼0.75-4.5Hz; δ) power during non-rapid-eye-movement sleep (NREMS), also termed slow-wave activity (SWA), is in a quantitative and predictive relationship with prior sleep-wake history, such that with increasing wake duration, there is a proportional increase in the level of δ power during early NREMS (Dijk et al., 1987, Franken et al., 2001, Brunner et al., 1990, Feinberg et al., 1992, Jones et al., 2008, Martinez-Gonzalez et al., 2008). Modeling approaches have established that most of the variance in δ power can be explained by the sleep-wake distribution, through the assumption of exponential saturating functions, which estimate increases in sleep pressure during waking, and continuous and gradual decreases across consecutive NREMS episodes (Achermann and Borbely, 1990, Franken et al., 1991, Franken et al., 2001, Guillaumin et al., 2018, Vassalli and Franken, 2017).

The predictability and accessibility of EEG δ power as a sleep-need marker has led to its widespread use in the field, and its inherent dynamics are now often equated with the sleep homeostatic process it is thought to reflect. In addition, several influential hypotheses on sleep regulation and function are based on how fluctuations in this variable reflect homeostatic sleep need (Daan et al., 1984, Krueger et al., 2008, Tononi and Cirelli, 2006). EEG δ power is quantified through frequency-domain analysis, using a Fast Fourier transformation (Borbely, 1982) and merges amplitude and incidence information of slow-waves (SWs) within the specified frequency range into a single metric. Time-domain analysis, quantifying other aspects SWs, have yielded new insights into their functional significance (Carrier et al., 2011, Freyburger et al., 2017, Vyazovskiy et al., 2009, Massart et al., 2014). For example, increases and decreases in SW slopes and changes to waveform profiles, are believed to reflect a recalibration of neuronal connections that strengthen during preceding waking (Fattinger et al., 2014, Panagiotou et al., 2017, Vyazovskiy et al., 2011, Tononi and Cirelli, 2006).

SWs are the defining electrophysiological feature of NREMS. The SW-frequency range comprises the slow-oscillation(SO), as well as δ oscillations (Neske, 2015, Steriade, 2006) with the latter being especially prevalent during NREMS after extended periods of wakefulness (Borbely et al., 1981, Feinberg et al., 1988). Although the term SW has been Although the term SW has been often used to solely describe the SO, here we use it to refer to both the SO and δ oscillations. In our use of the term SW, we refer to both the slow-oscillation and δ oscillations. It is still unclear how the various aspects of NREMS SWs respond to sleep-wake history and what their neuro-anatomical substrate and function is. Through detailed analysis of what causes the SW-period shortening we observed following prolonged periods of waking, we found that the well-known δ power changes are in fact a composite of the contributions of two separate SW populations each with profoundly different sleep-wake driven dynamics. Specifically, only SWs of the faster population were sensitive to prior time-spent-awake and returned to baseline levels much quicker than slower ones, findings which were similarly observed in humans. Furthermore, optogenetic silencing of the centromedial thalamus (CMT) in mice, increased only activity of this faster SW population, indicating that besides a dynamic distinction, there is an anatomical difference in their source of origin. Finally, the dynamics of the faster SWs were paralleled by by several physiological markers in both species. This study identifies a previously unknown complexity of the central and peripheral processes associated with the aftereffects of prolonged waking, typified by the short-lived decay of a specific sub-population of SWs, before reverting to levels typical of NREMS. This presents an inherent paradox, as the deepest levels of NREMS when recovery is presumed to be most efficient, in fact display several signatures more reminiscent of waking.

## Results

### NREMS slow-waves accelerate after prolonged periods of wakefulness

We first examined the effects of cortical surface recording site on sleep-wake driven dynamics in various features of EEG SWs during NREMS. Frontal (F)-, central (C)-, and, parietal (P)– placed electrodes were referenced to the cerebellum (Ref). The resulting differential signals were subjected to analyses in both the time-domain, to extract SW-amplitude, -slope, and -period of individual SWs (Fig. S1 A-D), and in the frequency-domain, to quantify δ (0.75-4.5 Hz) power [**see Methods**]. Time spent in NREMS during baseline showed a time-course typical of C57BL/6J male mice with the highest and most stable levels during the 12h light period, lowest in the first 6h of the dark period, and intermediate levels in the last 6h (Franken et al., 1999) (Fig. 1A **bottom**). A 6h SD resulted in NREMS increases during subsequent recovery, especially noticeable in the dark period (2-way rANOVA: interaction factors SD x time; *F*_17,170_ = 2.6: p=0.0009). Following prolonged periods predominated by waking (beginning of the baseline dark and during SD) the highest levels of δ power were observed, whilst during times when NREMS prevails; e.g., light phase during baseline and after SD, it was decreased to stable low levels reached in the last 4h of the baseline light periods. Although all 3 cortical derivations displayed this pattern, the highest relative power reached in both baseline and following SD, was seen in the frontal derivation (F-Ref), as compared to the other two (C-Ref, P-Ref; Fig. 1A **top**; 2-way rANOVA interaction factors derivation x time; *F*_26,390_ = 2.92: p<0.00001), previously reported in humans and rodents (Cajochen et al., 1999, Huber et al., 2000, Vyazovskiy and Tobler, 2005). Many studies in mice, including our own, rely on a bipolar F-P signal, which offers the clearest distinction of sleep-wake states (Mang and Franken, 2012). While absolute δ power levels during the baseline reference period (Fig. S2A-D) were higher in the F-Ref than the C- or P-Ref derivations, F-P and F-C levels and dynamics following SD were remarkably similar throughout the experiment, (Fig. 1A**-middle**). These latter findings imply that an electrode placed on the frontal cortical surface is sufficient for capturing maximal sleep-wake driven dynamic changes in the δ power range, regardless of the position of its bipolar pair.

**Figure 1:**
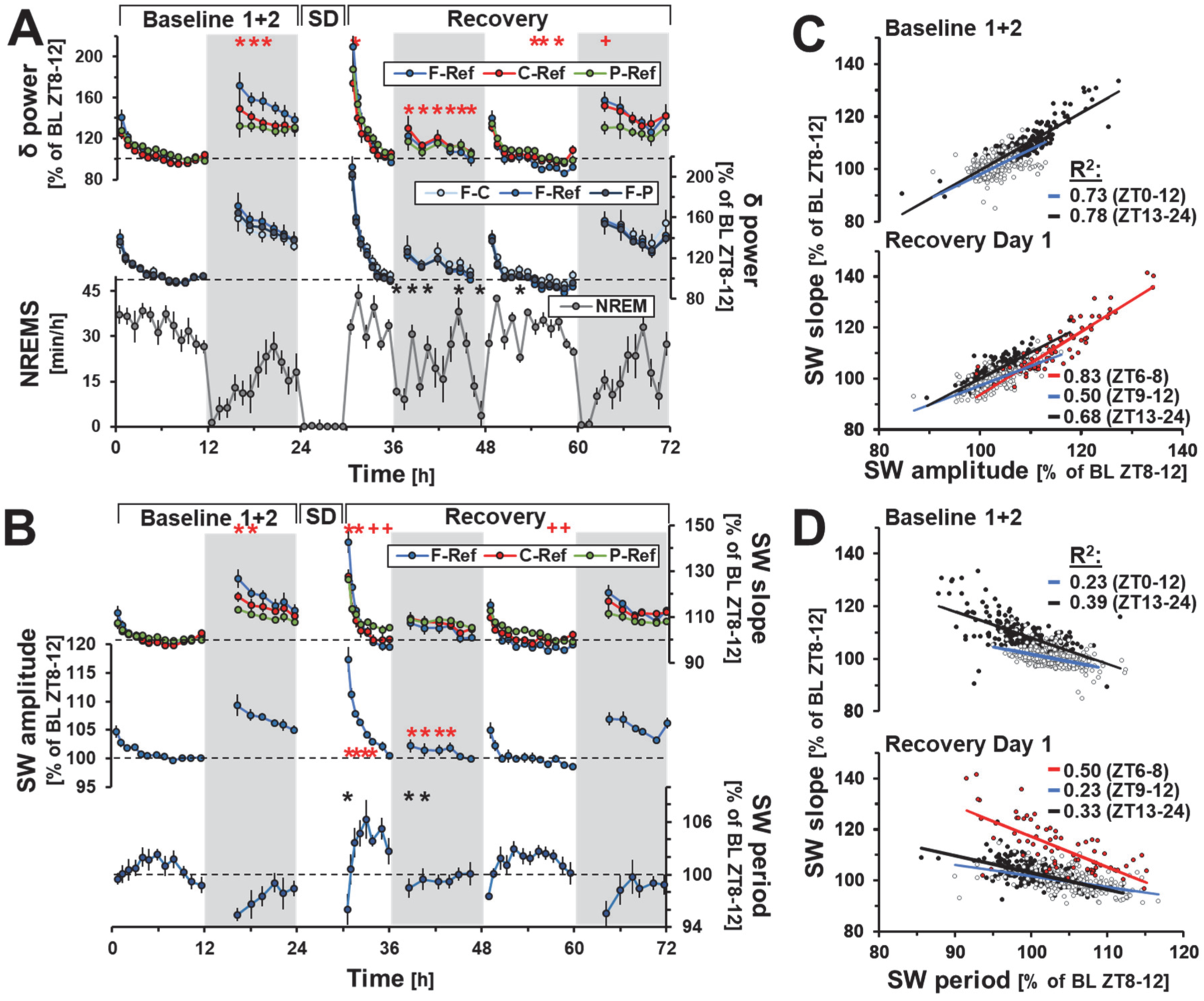
Topographical characteristics of time and frequency analysis of NREMS SWs before and after sleep deprivation (SD) **(A)** Relative increases in δ power consecutive to SD (24-30h) were site-specific and highest in frontal areas (**top**), independent of bipolar derivation (**middle**). Time-course of NREMS per hour of recording (**bottom**). **(B)** SW-slope changes were similar; however dynamic range was smaller (**top**). SW-amplitude (F-Ref) followed a nearly identical pattern (**middle**), whereas SW-period differed and was at times in opposition (**bottom**). Values represent mean +/- S.E.M (n=6) referenced to the baseline period of lowest δ power (ZT8-12). Red asterisks denote significant post-hoc differences (Tukey) between F- and C-/P-Ref derivation, and red plusses between C- and P-Ref (p<0.05). Baseline values are presented as an average across the 2 days preceding SD. (**C**) Linear regressions between SW time-domain components, during baseline and 18-hours of SD recovery in F-P derivations. SW-amplitude and -slope (**top-left**) were strongly correlated under baseline light and dark periods (grey and black circles, respectively), further increasing during the first third of SD recovery light period (ZT6-8; red circles). (**D**) SW-period showed higher correlations to SW-slope during dark periods, though was increased during the first third of the SD recovery light period. Lines represent linear regressions during respective time periods. Values represent mean values +/- S.E.M (n=38) referenced to baseline (ZT8-12). All linear regressions in **C** and **D** were statistically significant (p<0.05). For further details see Figs. S1/2.

Time-course analyses of SW-slope and -amplitude, revealed similar sleep-wake driven dynamics as δ-power, yet their range was smaller (Fig. 1B**-top** and **middle**, respectively; Fig. S1). Both SW-slope and δ-power mathematically depend on SW-amplitude, however spectral power scales as a square to this value whereas SW-slope scales linearly, exemplified by the fact that square-rooting raw δ power reduces its dynamic range to that of SW-slope (Fig. S2F). The largest SW-slope and -amplitude dynamics were again seen in frontal brain regions and were significantly higher than others (2-way rANOVA interaction factors: derivation x time; Amplitude-*F*_13,390_ = 34.3: p<0.00001, Slope-*F*_13,390_ = 35.9: p<0.00001; Figs. 1B, S2A-E). The similarity of the sleep-wake driven dynamics in SW-slope and -amplitude was substantiated by high correlations both during baseline [R^2^=0.849, n=684 percentiles (38 mice of 18 percentiles), p<0.000001], and SD recovery [R^2^=0.874, n=532 percentiles, p<0.000001; linear regression of all NREMS percentiles, including 32 additional mice from previous studies (Diessler et al., 2018, Hoekstra et al., 2019) with frontal-parietal derivation only; Figs. 1C,D; S2G)]. δ power was also strongly correlated to both SW-amplitude and -slope during baseline (R^2^=0.815 and 0.832, respectively, n=684 percentiles, p<0.000001) and following sleep deprivation (R^2^=0.845 and 0.837, n=532 percentiles, p<0.000001). This was not the case with SW-period, as correlations explained substantially less of the variance (baseline: R^2^=0.182, recovery: R^2^=0.108, p<0.000001), and differed statistically from amplitude/slope correlations (t-test on Fisher’s Z-transformed r value; baseline: p<0.00001; recovery: p<0.00001). In addition, the lowest values for SW-period were present specifically within the first hour of recovery (red circles), where relative increases in SW-slope and -amplitude were highest, as compared to the remainder of the light or dark period (white and black circles, respectively). Thus, much of the observed variance in SW-slope was due to these initial values of SW-period at recovery onset as the former largely co-varies with SW-amplitude, except when sleep pressure is high (i.e., during the first NREMS episodes during the dark phase and immediately following SD, Fig. 1B,D**-bottom,** Fig. S2G, H). Finally, this shortening of SW-period did not significantly differ among cortical sites (Fig. S2E**, right**).

### Time dynamics of NREMS δ-band frequencies are not homogenous

Separating the first 6-hours of SD recovery into 25 percentiles each containing approximately the same number of minutes of NREMS from consolidated (>32s) episodes (mean NREMS per percentile = 7.0 ± 0.1m), we focused on the time period where the largest changes in SW-period were observed (≈ first 2 hours, 8 percentiles). We found that the highest initial amplitude and quickest subsequent decreases were for individual SWs with a period between 0.25-0.5s (i.e., 2-4 Hz) (Figs. 2A, S3A). In addition, the incidence of these faster SWs was affected the most at this time (Fig. S3B). Importantly, we discovered a bi-modality in the prevalence of SWs with peaks centered at 1.25 and 3 Hz that became even more striking when corrected for the inherent frequency bias (Fig. 2B), as slower SWs cannot physically occur as often as those which are faster. Correlations between amplitude and slope by frequency during this time where also highest (R^2^>0.9) above 2Hz (Fig. 2C), a finding previously noted in humans following SD, albeit at slower frequencies (Bersagliere and Achermann, 2010). Evidence of frequency-specific dynamics was also obtained (Fig. S3G). Estimating the timepoint during SD recovery at which each 0.25-Hz bin had lost half of its initial power (t_1/2_; Fig. S3E), we found that faster δ band frequencies (2-4.5 Hz) showed steeper declines, in contrast to slower ones (0.5-1.75 Hz). Notably, frequencies above the δ range also decreased more quickly than these slower waves, however their initial values were much lower (Fig. S3F). Examination of changes to the power spectra across time confirmed that the largest initial increases were confined to these faster oscillations and indeed the majority of the spectrum, displaying highly significant differences between the 1^st^/2^nd^ (2-way rANOVA: percentile x frequency; *F*_98,7252_ = 9.37: p<0.000001) and 2^nd^/3^rd^ percentiles (*F*_98,7252_ = 2.28: p<0.00001), though not after (3^rd^/4^th^: p=0.35; Fig. 2D). To identify where these deviations were greatest, we analyzed percentile-to-percentile differences using moving averages of 5 consecutive 0.25-Hz bins and found the largest decreases to occur between 2.5-4.5 Hz, while, surprisingly, power in the slowest frequencies (centered at 0.5-1.0 Hz) initially increased (Fig. 2E; 2-way rANOVA: percentile (slope) x frequency; *F*_99,7326_ = 9.13: p<0.00001). Finally, using an unbiased hierarchical clustering approach on the time course of EEG power density at 0.25-Hz resolution during these first 8 percentiles we identified a lower-(0.5-1.75 Hz) and higher-frequency δ band (2-4.5 Hz; Fig. S3C). The aggregate results of these SW time-course approaches (Figs. 2E, S3C, S3E) and SW incidence distribution (Fig. 2B), strongly indicated two distinct δ sub-bands separated at around 2.0-2.25 Hz. As the demarcation of these two bands did not appear static, we chose 5 representative 0.25Hz bins for each based on this separation (δ1: 0.75-1.75; δ2: 2.5-3.5 Hz).

**Figure 2:**
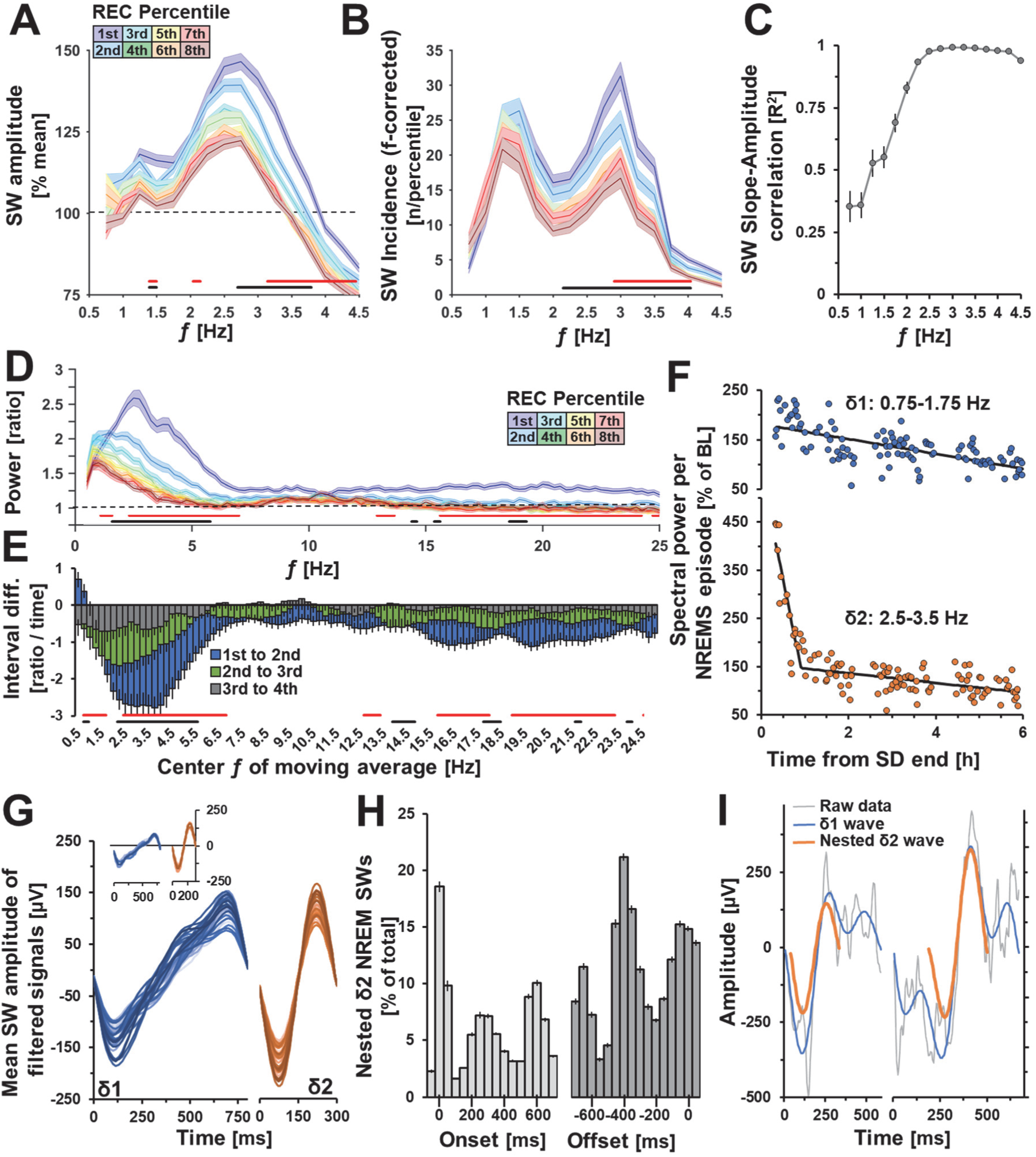
Evidence for two distinct δ bands. Distribution of **(A)** SW-amplitude and **(B)** SW-incidence according to SW period (frequency; 0.25Hz bins) during the first 2h of SD recovery (8 percentiles). SW incidence, corrected for frequency (n/frequency), shows a bimodal distribution. (**C**) SW-slope-amplitude correlations during the same time were highest in faster frequencies. (**D**) Relative spectral power during the first 2h of SD recovery, displays not only a significant decrease in power, but a shift from faster to slower frequencies. (**E**) Ratios of five 0.25 Hz bin moving averages between spectra in the 1^st^-2^nd^, 2^nd^-3^rd^, and 3^rd^-4^th^ time percentiles in D. (**F**) Time-course of δ1 and δ2 power during the first 6h of SD recovery per NREMS episode in one mouse. (**G**) Average waveforms of detected δ1 (**blue**) and δ2 (**orange**) waves during the first 10 minutes of recovery NREMS following SD, for individual mice (n=38). Note the choice of using a fronto-cerebellar derivation does not affect these waveforms (**insert**; n=6). (**H**) Timing of nested δ2 waves relative to δ1 onset (**left**) or offset (**right**; = 0ms for both). Histograms span 50ms time bins plotted at midpoint. Of note, the second and third histogram peaks indicate the starts of these additional nested δ2 waves. (**I**) Example of a δ2 wave (**orange**) at onset (**left**) or during (**right**) a δ1 wave (**blue**). Unfiltered signals are presented in grey. Error bars and shaded areas are ± S.E.M. Significant changes between 1st-2nd and 2nd-3rd percentiles are indicated with red and black bars, respectively (n=38). For further details see **Fig. S3/4**.

Reanalysis of SD recovery dynamics of these two δ bands, within individual consolidated NREMS episodes, showed that mean initial values (first percentile) were significantly different (δ1: 174 ± 6%, δ2: 248 ± 11%; t-test p<0.00001). Furthermore, recovery dynamics during the first 6 hours differed between the two, with δ1 increasing and subsequently decreasing linearly, and δ2 falling rapidly within the first hour, followed by a more gradual decline (example in one mouse, Fig. 2F). To estimate the point in time at which the rate of decrease in δ2 changed (“pivot point”), dynamics were subjected to a two-segment piecewise linear regression (black line), which was present in all mice (59.7 ± 7.2 min after NREMS onset, see Fig. S4). The decay in δ1 did not differ before and after pivot point (t-test p=0.55) and was much slower than δ2 [δ1 = −35 ± 14; δ2 = −166 ± 12; reference power (%)/hour; t-test p<0.00001]. Importantly, the δ2 decay after the pivot point was similar to δ1 (δ1 = −26 ± 4; δ2 = −20 ± 3 %/hour; t-test p=0.24). Finally, as the pivot point varied across animals, as well as the decay rate for δ2, averaging gave the appearance of an exponential decay, which was not reflected in individuals (Fig. 2F, S4).

### δ-2 SWs are nested inside of δ-1 SWs

With these frequency definitions, we reanalyzed the mean waveforms of δ1 and δ2 SWs during initial NREMS episodes after SD. δ1 SWs (Fig. 2G**; left**) resembled what has been previously described in rodents and humans as a multipeak SW (Panagiotou et al., 2017, Vyazovskiy et al., 2007, Riedner et al., 2007). Conversely, the profile of δ2 SWs was uniformly sinusoidal (**right**; Fig. 2G). These overall waveforms persisted when the raw signal was analyzed (Fig. S3D) and did not depend on derivation (**see insert: F-Ref**). To further explore the composition of δ1 SWs, raw signals were filtered along a narrower band encompassing only δ2 frequencies (2-4.5 Hz). Surprisingly, we found that all δ1 SWs contained nested δ2 oscillations (96.6 ± 0.6%), and of those, more than half contained at least two (65.4 ± 2.0%), representing a quarter (25 ±0.4%) of all δ2 waves across the first 6h of recovery. When we determined their nested position, we identified a predominance for them either to start (Fig. 2H, I**; left**) or end (Fig. 2H**, right**) with the δ1 SW, and at times occur in the center (Fig. 2I**; right**). Importantly, these data were not randomly distributed (onset; 1-way rANOVA: bin; *F*_15,540_ = 305.2: p<0.00001; offset; 1-way rANOVA: bin; *F*_15,540_ = 192.8: p<0.00001).

### δ-2 power, but not δ-1 power is in a quantitative relationship with prior sleep-wake history

Given the radical differences observed in the dynamics of these two bands during the first three percentiles (as in Fig. 2D; 24.8 ± 0.9min NREMS), we probed this time-period under a variety of conditions. At wake-to-NREMS transitions immediately following recovery (8.9 ± 0.5min NREMS), δ2 significantly increased as compared to δ1 (two-way rANOVA: sub-band x time *F*_44,352_ = 4.8, p<0.000001), rising to its maximum levels after 32s-64s (Fig. 3A; **left**). δ1 also increased, though calculated rise-to-maximum regressions indicated a 4-fold slower buildup (slope-to-maximum: δ1=48.6, δ2=228.6; reference power %/min.). Differences between the bands dissipated afterwards (**right**).

**Figure 3:**
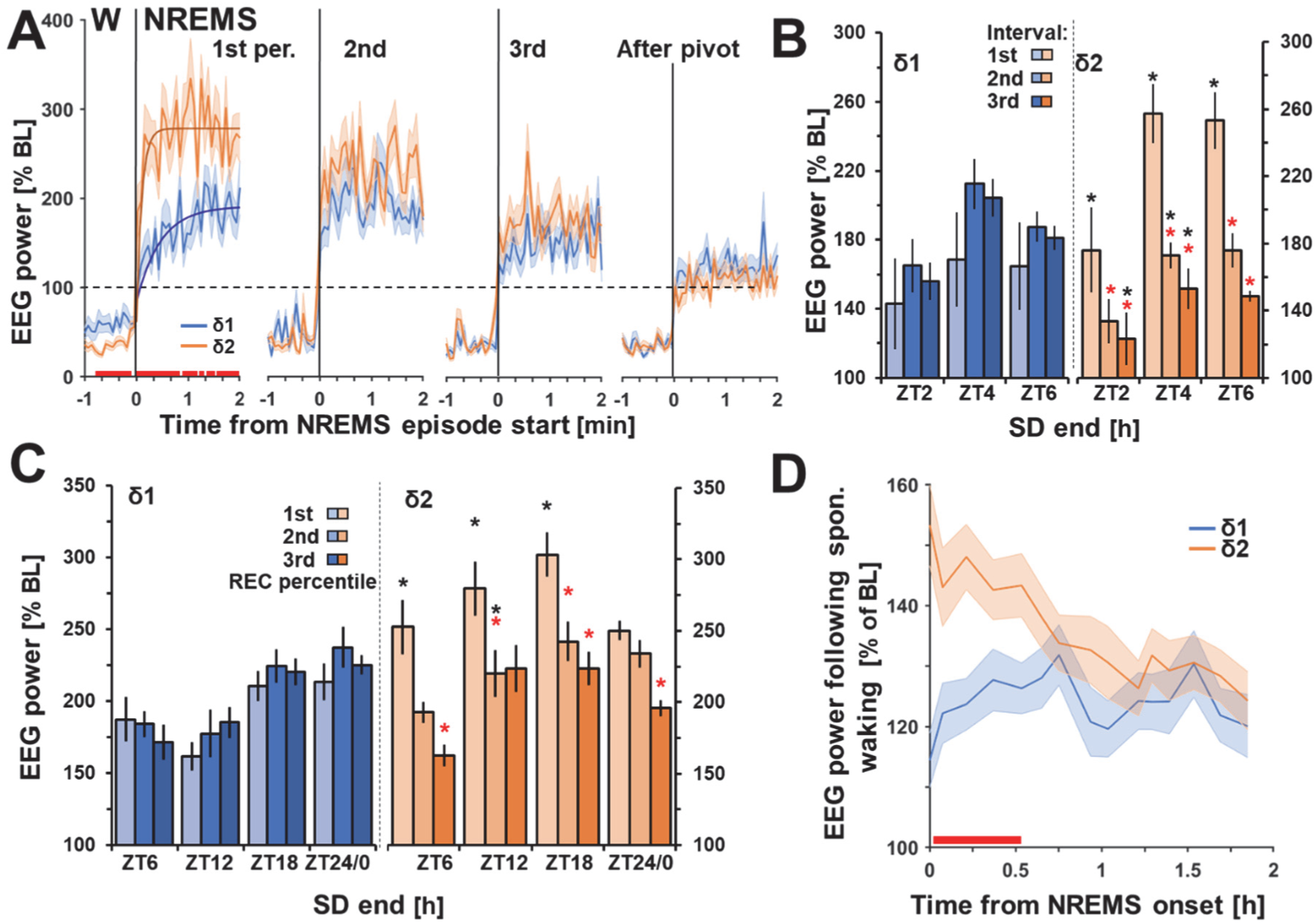
δ1 and δ2 behave differently following enforced and spontaneous extended waking. (**A**) δ1 and δ2 across wake (W) to NREMS transitions during recovery from 6h SD. Lines represent ‘rise-to-maximum’ regressions. (**B**) SDs starting at light onset (ZT0) of different lengths (2, 4, 6h) (n=7/group). Asterisks represent Tukey post-hoc significance (p<0.05) between first percentile (red), or δ1 vs. δ2 (black). (**C**) Effect of time-of-day at which SD ends on δ1 and δ2. Asterisks same as in **B** (n=8/group). (**D**) NREMS following prolonged spontaneous waking (1.6 ± 0.1h) during the baseline dark period showed initial increases for δ2 power which then decreased across subsequent NREMS episodes, whereas δ1 did not. Red lines same as asterisks in **B, C** (p<0.05; n=38). Data is presented as mean values, error bars and shaded areas are ± SEM.

Next, we quantified the δ1 and δ2 dynamics in the three first percentiles recovery following SD of varying lengths (2, 4, and 6h). We found that δ1 power was unaffected time spent in SD (Fig. 3B**, left**), while a significant increase was observed from the 1^st^ to 2^nd^ recovery percentile (1-way rANOVA: time; *F*_2,36_ = 9.2 p=0.0006). In contrast, δ2 showed immediately higher power, followed by rapid decline, which was already seen after 2h of SD (2-way rANOVA: SD length x percentile; *F*_4,36_ = 4.3, p=0.006), and at higher levels than δ1, which was not significant over time (2-way rANOVA: SD length x percentile; *F*_4,36_ = 0.7, p=0.62). Thus, only δ2 seemed to be in a quantitative relationship with prior time-spent-awake.

To assess whether time-of-day affected recovery dynamics, we reanalyzed a previously published dataset, where 6h SDs were performed starting at either ZT0, 6, 12, or 18 (Curie et al., 2013). Initial values of δ1 and δ2 varied with time-of-day but only δ1, significantly [1-way rANOVA: SD end; *F*_3,27_ = 4.12, p=0.02; Fig. 3C], with the highest levels after SD end at ZT24/0. In δ2, this was seen at ZT18, where the persistence of waking was longest (NREMS onset after SD end: 2.8 ± 0.2h; i.e., ∼9h of waking). SD end time did not affect δ1 dynamics (2-way rANOVA: SD x time; *F*_6,54_ = 1.44, p=0.3), whereas δ2 dynamics were significantly different (2-way rANOVA: SD x time; *F*_6,54_ = 2.48, p=0.03). This seems to suggest that while δ2 is sensitive to prior time-spent-awake, initial increases in δ1 may be sensitive to time-of-day. In a forced desynchrony protocol in humans (Lazar et al., 2015), SW-period was longest during the subjective dark period (approx. 2AM), perhaps due increases in the incidence of slow δ oscillations as a function of circadian time. We found that initial values of δ1 were highest during recovery at the beginning of the light period, which may reflect something similar.

We next considered whether spontaneous prolonged waking bouts similarly provoked these differential dynamics. Baseline dark periods began on average with extended periods of waking (1.6 ± 0.1h) followed by consolidated NREMS, a time at which we had previously observed shortening of the SW-period (Fig. 1B). Power in both δ sub-bands were increased above baseline levels (% of ZT8-12) although the increase for δ2 was significantly higher than δ1 (2-way rANOVA: δ sub-band x time; *F_14,1036_* = 4.14: p<0.00001; Fig. 3D). Surprisingly although δ1 was above baseline reference values, a lack of decay across time was observed. Through these experiments, we found that dynamics for both δ1 and δ2, as well as the differences between them, were consistent and robust. Although δ1 always increased after SD, this was smaller than δ2 and importantly, was not in a quantitative relationship with prior waking and did not change over subsequent NREMS percentiles, which are defining criteria for homeostatically regulated variables.

### Typical δ-2 dynamics are observed throughout the thalamus and cortex, but are most pronounced in frontal areas

The previous observation of the frontal dominance of δ power following prolonged waking (Fig. 1A) was found to be due solely to δ2, while δ1 dynamics did not vary across cortical surface (Fig. 4A). As the greatest differences between electrode sites were identified during the first percentile of SD recovery (7.7±1.4m NREMS, 23.6±1.3min total), we compared power spectra at this time to ascertain frequency specificity of this effect (Fig. 4B). Power spectra were significantly different from one another depending on cortical site (2-way rANOVA: site x frequency bin; *F_196,1176_* = 2.38: p<0.00001), with the largest differences seen in the δ2 range, especially between frontal and parietal electrodes. Furthermore, while there was no difference between frontal and central areas at frequencies above δ (10-25 Hz), both were significantly higher than the parietal (Fig 4B; dark/light grey lines).

**Figure 4:**
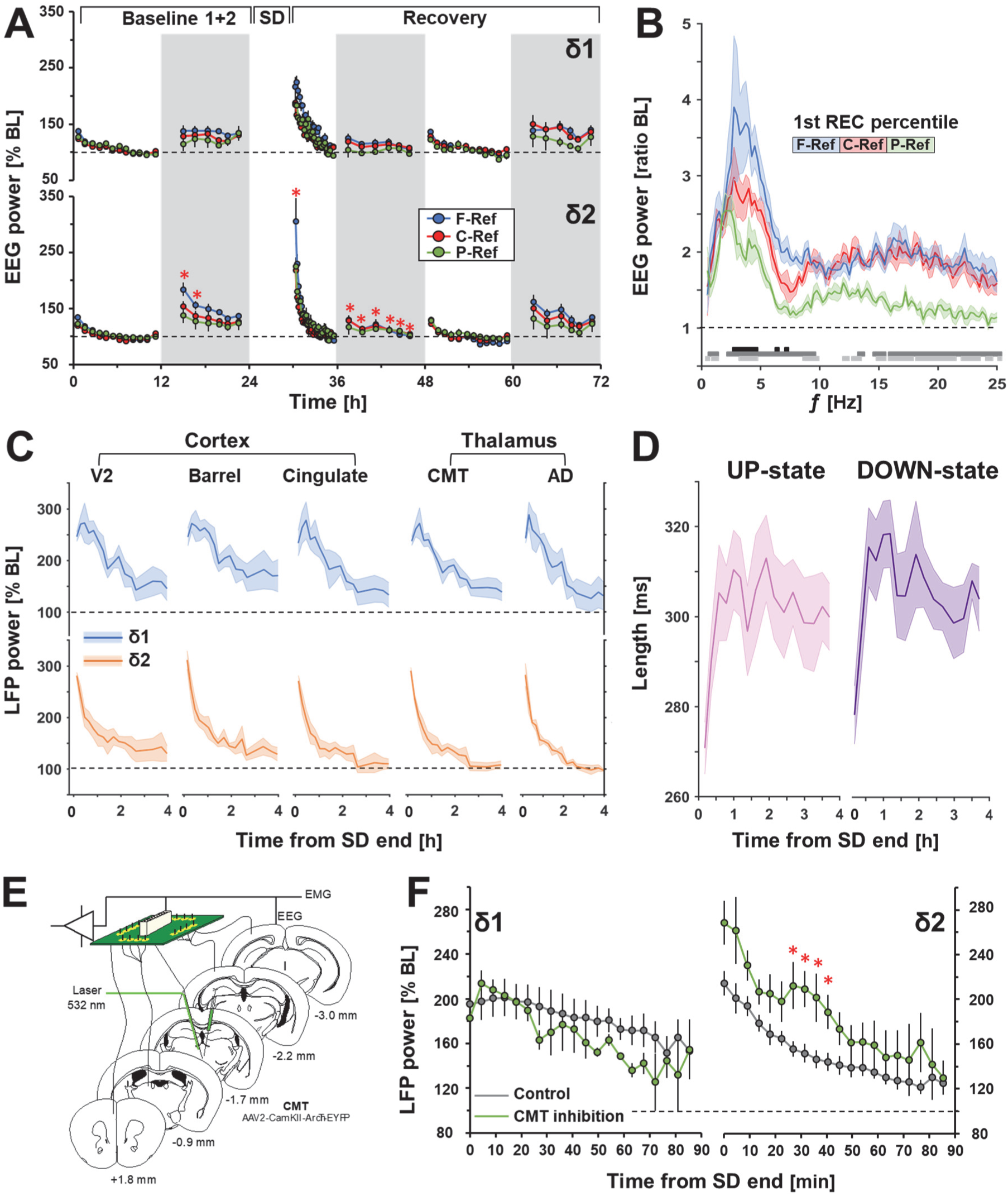
δ2 dynamics are visible throughout the thalamus and cortex and are altered following thalamic silencing. (**A**) Time-course of δ1 and δ2 at different cortical surface areas. δ2 increases at specific times during baseline (dark period) and following SD, and was highest in frontal areas, whilst derivation did not affect δ1. Red stars denote post-hoc significance (Tukey) between F-Ref and C/P-Ref. (**B**) Power spectral density (PSD) across recording site during initial recovery NREMS (first 5 minutes). Black, dark grey, and light grey lines represent post-hoc significant differences (p<0.05; Tukey) between Fr-Ce, Fr-Pa, and Ce-Pa, respectively (n=6). (**C**) LFP recordings following SD (4h) in multiple brain structures showed similar dynamic changes in both δ1 and δ2 as **A**. (**D**) Average length of UP (left) and DOWN (right) states at the same time in the cingulate cortex during NREMS. (**E**) Optogenetic inhibition of the CMT increased δ2 power in NREMS (**F; right**) in the cingulate cortex following SD, as compared to control animals (**grey**). Red asterisks denote significant differences (p<0.05; Tukey) from control. No effect is seen on δ1 dynamics (**left**). Error bars in **A**, **D**, **F** and shading in **B**, **C** indicate ± S.E.M; **A, B** n=6 mice, **C-F**, n=4 mice per group.

To gain insight into the underlying circuitry, we probed deeper cortical and sub-cortical layers (thalamus), using a published dataset (Gent et al., 2018a) of local field potential (LFP) tetrode recordings at 3 cortical [Visual (V2), Barrel, and Cingulate cortices] and 2 thalamic sites [centromedial thalamus (CMT), anterodorsal nucleus (AD)], at varying depths. Following a 4h SD ending at ZT4, high levels in both δ1- and δ2-power were observed, in visually similar patterns to cortical surface electrodes and strikingly similar across recording sites (Fig. 4C). δ2 displayed a rapid decrease in power across all sites, and δ1 an initial increase before a linear decrease. Finally, we found that during SD recovery, UP/DOWN state lengths recorded from the cingulate cortex, were initially shorter and returned to stable levels as sleep progressed (rANOVA time: *F_15,_*_90_=11.735: p<0.00001; Fig. 4D), reminiscent of the shortening of SW-period observed after SD (Fig. 1B). Although the frequency of the number of UP and DOWN states was higher after SD, average spiking rates across recovery did not change and was consistently greater for UP- vs. DOWN-states (13 ± 3 and 4 ± 1 Hz, respectively).

The CMT is thought to be a critical component of the circuit underlying the generation of brain-wide SWs during NREMS (Gent et al., 2018b). Optogenetic silencing of CMT neurons (Fig. 4E) was previously shown to increase NREMS δ power during SD recovery sleep in the cingulate cortex, which receives direct input from the CMT (Gent et al., 2018a). We found that during CMT inhibition this increase was specific to the δ2 band, leaving δ1 activity unaffected (Fig. 4F).

### Distinct δ sub-bands are present in the human brain following sleep deprivation

We next assessed whether humans showed a similar heterogeneity in the response of δ power to SD. A total of 110 healthy human subjects from four published datasets who underwent whole-night SD amounting a sustained waking period of 40h (Bodenmann et al., 2009, Holst et al., 2017, Valomon et al., 2018, Weigend et al., 2019), were included in the analysis. Here we show only the aggregate results, but similar results were obtained in each study separately. Using a similar approach as with mice and focused on the right fronto-parietal (F4-P4) derivation (compare Fig. 5A to Fig. 2E), we identified clear separations in frequency bands between consecutive NREMS episodes. Of note, other derivations assessed in the frontal areas (i.e. Fz, left F3-P3) displayed similar relative changes (data not shown). As in mice, changes in humans within the δ band appeared heterogenous, with lower frequencies initially increasing and faster ones decreasing (2-way rANOVA: percentile-percentile difference x frequency; *F_98,12544_* = 11.6: p<0.00001). Using these results, we took as representative bands for δ1 and δ2, 0.5-1.0Hz and 1.5-2.0Hz, respectively. After SD, δ2 was initially high and decreased rapidly across the first two NREMS episodes (δ2; 2-way rANOVA: SD condition x time-course; *F_49,8134_* = 29.4: p<0.00001), as compared to δ1 which was slightly increased in second NREMS episode vs. the first (δ1; 2-way rANOVA: SD condition x time-course; *F_49,8134_* = 7.26: p<0.00001; Fig. 5B**, bottom;** 2-way rANOVA: δ1/2 x time-course; *F_49,8134_* = 23.7: p<0.00001). After 16h spontaneous waking (baseline), δ1 exhibited a slower decline than δ2, which was lower during the second NREMS episode, similar to what we previously found in mice (**top**; 2-way rANOVA: δ 1/2 x time-course; *F_49,8134_* = 3.59: p<0.00001). During baseline, δ2 was also higher in the first NREMS episode, though lower than after SD (Fig. 5B**, top**). Maximum values during the first and second NREMS episodes for baseline and SD recovery showed both higher initial values and steeper decays for δ2 than δ1 (Fig. 5B**, insert**).

**Figure 5:**
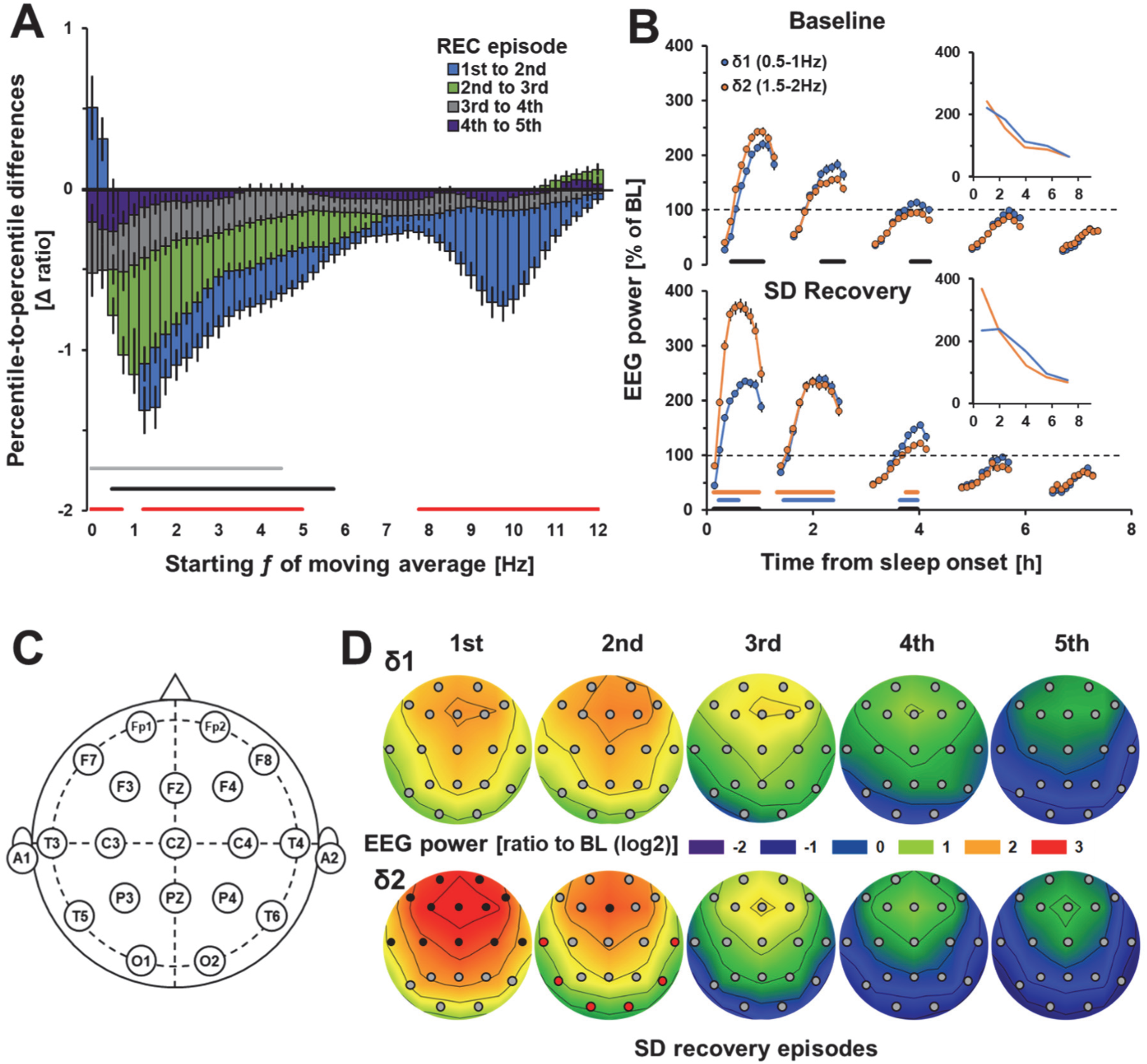
Recovery following SD reveals a δ-band heterogeneity in humans. (**A**) Differences in moving averages of three consecutive 0.25Hz bins of maximum peaks in the right frontal-parietal derivation (F4-P4) of NREMS episodes following a 40h sleep deprivation during the 1^st^-2^nd^ (blue), 2^nd^-3^rd^ (green), 3^rd^-4^th^ (grey), and 4^th^-5^th^ (dark blue). The largest decrease was observed in faster δ (1.5-2.0Hz), while in slower δ (0.5-1 Hz) frequencies were initially increased. Red and black lines represent post-hoc significance (p<0.05), 2^nd^/1^st^, 3^rd^/-2^nd^, and 4^th^/3^rd^, respectively. (**B**) Time-course plots of δ1 (0.5-1.0Hz) and δ2 (1.5-2.0Hz) power during each NREMS episode (10 percentiles/episode) in undisturbed conditions (baseline-**top**) and following SD (recovery-**bottom**), gaps reflect intervening wake or REMS. After SD, δ2 was increased during the first NREMS episode, illustrated by the time-course of maximum values per episode (**insert**). Black lines represent post-hoc significance (Tukey) between percentiles of δ1 and δ2, blue and orange between δ1 and δ2 from baseline to recovery, respectively. Values are given as percentage of baseline mean ± S.E.M (black lines) (n=110). (**C**, **D**) Electrode placement, and heat plots in a subset of healthy volunteers (n=21) with 19 electrode sites, of δ1 and δ2 during SD recovery. Increases in δ2 were highest in frontal areas and dissipated more quickly in subsequent episodes than δ1. Values are expressed as log2 of ratio to mean power across all electrode sites during BL, for each subject. Black and red filled circles represent statistically different electrode sites where δ2 was higher and lower, respectively, than δ1 (Tukey post-hoc p<0.05). Black contour lines are fixed at 0.5.

To determine whether dynamics were region-specific or represented a global process across the cortex, a subset of individuals (n=21) with 19 recording sites were analyzed [Fig. 5C; (Weigend et al., 2019)]. After SD, the highest relative power increases for δ2 were confined to frontal areas (Fig. 5D). Interestingly, only the first NREMS episode showed significant differences between the two bands across more than half of the surface. During the second NREMS episode some significance persisted in frontal areas, though δ2 decreased faster in more posterior areas falling below δ1 relative levels (red circles). Our results demonstrate that differential dynamics of δ1- and δ2-power are present in humans, although both δ bands center at lower frequencies than in mice.

### δ2 dynamics typify the time-course of physiological changes during NREMS after prolonged waking

The short-lived nature of high δ2 levels after SD was evidenced in mice where nearly half of the initial power was lost during the first hour of recovery (δ2_initial_ = 249 ± 11%, δ21h = 136 ± 3% of reference) and between NREMS episodes 1 and 2 in humans (δ2_1st_episode_ = 369 ± 12%, δ2_2nd_episode_ = 233 ± 9%; Figs. 2F, 5B; Fig. S5D). Given this observation, is sleep homeostasis as evidenced by the dynamics of δ power truly a continuous process, or does the non-continuous rapid decay in its main component point to a discrete sub-state of NREMS, emerging only after prolonged periods of wakefulness? During this early recovery phase, a number of additional variables reached values atypical of NREMS that all reverted to baseline levels before pivot point. We already noted that spectral power of higher EEG frequencies during NREMS (beta/low-gamma 18-45 Hz) was significantly increased during initial episodes of recovery (higher than at any point during baseline; Fig. 2D) and when these frequencies were analyzed identically to δ1- and δ2-power, a familiar pattern emerged (Fig. 6B). Even more surprisingly, other physiological measures followed similar dynamics. Muscle tone during NREMS was initially high, followed by a rapid decrease before reaching the stable baseline values after the pivot point (Fig. 6C). In addition, cortical temperature (T_cortex_), recorded in a subset of mice, showed a significant reduction up to the pivot point, losing more than 2°C in less than 60 minutes (Fig. 6D), as compared to the high sustained values during SD. However, changes in respiratory rate, which in the mouse is known to occur between 3-4 times per second [3-4Hz; (Friedman et al., 2004)], overlapping with the δ2 band (2.5-3.5 Hz), did not correspond with observations made with these other measurements during NREMS at this time (Fig 6E).

**Figure 6:**
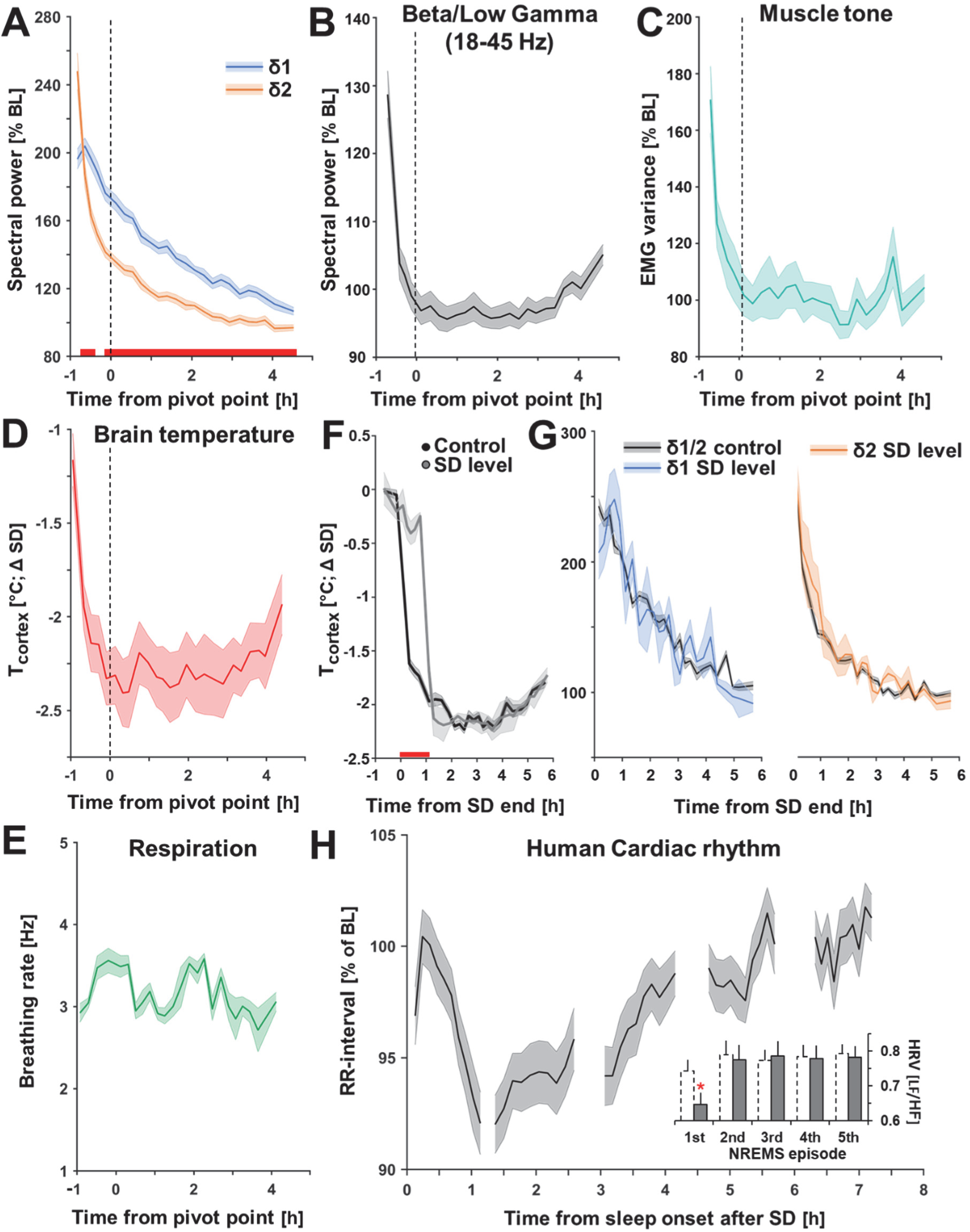
δ2 dynamics typify a physiologically different NREMS sub-state. **(A)** δ2 power (**orange**) decline across the first 6-hours of SD recovery compared to δ1 (**blue**). (**B**) EEG power in higher frequencies (beta/low-gamma: 18-45 Hz), (**C**) muscle tone, and (**D**) brain temperature, but not (**E**) respiratory rate. Of note, power reached in higher frequencies was greater following SD than at any point during baseline (**Fig. S5F**). (**F-G**) Maintaining cortical temperature at SD waking levels during the first hour of recovery in a subset of mice using a TEG (n=4), did not alter δ1 or δ2 dynamics. (**H**) Human cardiac rhythm (measured as R-R interval) differed during the first NREMS episode after SD compared others, also reflected in HRV (LF/HF ratio; insert). Dashed lines/grey histograms represent baseline and SD recovery, respectively. All values are mean (± S.E.M.) and red bars represent Tukey post-hoc significance (p<0.05). Vertical dashed lines in **A-E** represent average of individually calculated pivot points. **A-C, F** (n=38), **D** (n=8), **E** (n=3), **F, G** (n=4), **H** (n=40). For more detail, see **Figures S5/6**.

As we identified such marked changes following SD, we also sought to determine whether some of these variables were affected after periods of spontaneous waking under baseline conditions. NREMS muscle tone was also increased immediately after sleep onset during the dark periods (Fig. S5C), in conjunction with the incidence of δ2, but not δ1 waves, which were similar in dynamics to their respective power (Fig. S5A, B). This wake-history-dependent difference was further corroborated by analyzing δ2-δ1 power ratio, which saw the lowest values during the majority of the light periods, followed by increases in baseline dark and after SD (Fig. S5E). Examination of other frequency bands (theta, sigma, and beta/low gamma) during NREMS, showed similar behavior (Fig. S5F). In humans, we additionally observed changes between subsequent NREMS SD recovery episodes in faster frequencies (8-11Hz; Fig. 5A). Time-course analysis revealed significant relative increases (2-way rANOVA: condition x time-course; F_49,10535_ = 8.93, p<0.00001; post-hoc t-test) after SD only during the 1^st^ NREMS episode as well as raw spectral power (BL: 2.9 ± 0.2µV^2^, SD: 4.6 ± 0.4 µV^2^; t-test: p<0.00001), confirming changes in EEG signatures not directly related to δ2 activity (Fig. S5G), which has been previously noted (Dijk et al., 1990). In a subset these subjects (n=40) recorded with EKG, we observed that heart rate [HR; measured as % of mean BL R-R interval (1.07 ± 0.02s or 57 ± 1 beats-per-minute)], was significantly different from baseline and lowest in the first SD recovery NREMS episode [2-way rANOVA: condition x time-course; F_49,3822_ = 1.75, p=0.001; post-hoc Tukey t-test; (Figs. 6H, S5H)]. This initial difference was further reflected in heart rate variability (HRV; Figs. 6H, **insert**), as changes in the ratio between low-frequency (0.04-0.15Hz) and high-frequency components (0.15-0.4Hz) of HR, considered a measurement of sympathovagal tone (Eckberg, 1997). This is in line with previous studies that have noted other changes in physiological measures during the first NREMS episode, for example, increases in muscle tone (Brunner et al., 1990) and core body temperature (Dijk and Czeisler, 1995), giving credence to the idea of an active physiological state coinciding with the deepest sleep.

Changes in brain and body temperature have been shown to affect the frequency of EEG oscillations such as theta (6-9Hz) and the slow oscillation (<1 Hz) (Deboer, 1998, Sheroziya and Timofeev, 2015). To determine if temperature was causally related to δ2 dynamics, we maintained T_cortex_ at SD levels during the first hour of recovery using a thermoelectric generator (TEG). The TEG successfully sustained brain temperature at SD levels (Fig. S6, Fig. 6F) without affecting sleep-wake state (Fig. S6D). However, impeding the decrease of T_cortex_ did not affect the time-course of δ2 (or δ1)-power (Fig. 6G). Although increased temperature can speed up a particular oscillation, SD selectively enhances the production of δ2 waves more so than δ1, giving the impression of an overall increase in δ frequency. This set of physiological variables with comparable fast dynamics in the first hour for recovery (or NREMS episode in humans), provides evidence that early NREMS qualitatively differs from all subsequent sleep.

## Discussion

Through in-depth analyses of SW features in mice, we noted that during NREMS immediately following extended periods of waking, either spontaneous or experimentally evoked, δ-waves briefly increased in frequency. We discovered that this was not due to general shortening of SW-periods across the δ-band, but instead to an increased incidence and amplitude of SWs belonging to a population of faster δ waves (2.5-4.5Hz), which we termed δ2. Prevalence and amplitude of a slower population (0.75-2.0Hz; δ1) remained relatively unperturbed during this initial NREMS phase and preceding time-spent-awake did not predict their initial levels. The concomitant increase in amplitude and prevalence of δ2-waves fully accounted for the increase in SW-slope that we and others have observed after extended periods of wakefulness, and that has been interpreted as an independent SW-feature gauging homeostatic sleep pressure functionally related to synaptic strength (Tononi, 2009).

Perhaps the most salient result of our study is the short-lived nature of δ2-wave promotion, as within the first hour following SD, their features (power, amplitude, incidence) reverted to near baseline levels. These specific short-lasting and discontinuous dynamics (exemplified with the “pivot point”), are inconsistent with accepted models of sleep regulation and function based on those of a merged δ-band as a sleep-need proxy. δ2 recovery time-course was paralleled by an equally fast decay in the levels of a number of (neuro-)physiological variables not generally considered to be associated with the process of sleep homeostasis. We interpret this highly dynamic NREMS phase to reflect a transition during which the system adjusts from the aftereffects of a highly active and sustained waking period before typical NREMS can be reinstated.

### Two populations of SWs

The bimodal distribution of SW incidence across frequencies, the differences in the waveforms of δ1 and δ2 SWs, and their different dynamics in response to sleep-wake history, taken together argue for the existence of two classes of SWs. We are not the first to note a heterogeneity in the δ frequency range, but the frequency bands discerned were not always consistent (Amzica and Steriade, 1998, Deboer et al., 2002, Huber et al., 2000, Siclari et al., 2014, Vassalli and Franken, 2017). Our classification of the two δ sub-bands is based on differences in their sleep-wake driven dynamics *in vivo* and in unrestrained mice, therefore this division might yield alternative frequency boundaries as studies focusing on other aspects of SWs in different experimental contexts. Nevertheless, the implementation of three unbiased methods converged on ∼2.0/2.25Hz separating the δ1 and δ2 bands in the mouse, which matched remarkably well the results of other rodent studies (Deboer et al., 2002, Franken et al., 2006, Huber et al., 2000, Vassalli and Franken, 2017, Binder et al., 2012). Another important factor concerns species differences that have thus far been largely overlooked, as the same band is used (∼0.75-4.5Hz) to denote EEG δ-activity (SWA) in humans and rodents. With a similar approach to estimate the frequency separating the two δ-bands we arrived at ∼1.0/1.25Hz in humans; i.e., left-shifted by 1Hz compared to the mouse. Such inter-species frequency-slowing of oscillatory events, which have presumed common origins, was previously reported for the theta band and were suggested to relate to differences in brain size (Buzsaki et al., 2013, Watrous et al., 2013).

What we refer to as δ1 in humans falls in the frequency range generally reserved for the slow oscillation [SO; <1.0 Hz (Steriade et al., 1993)], the dynamics of which, similar to our findings, did not co-vary with time-spent-awake and asleep (Achermann and Borbely, 1997). Applied to the mouse, the SO (or δ1) upper demarcation might be as high as 2.5Hz, as others have already assumed due to other considerations (Binder et al., 2012, Fernandez et al., 2018). Accordingly, the δ1 population of SWs we identified, could represent the SO while δ2 SWs could represent δ, albeit with species-specific lower and upper frequency boundaries. However, in the remainder of the text we have kept our δ1/δ2 nomenclature when referring to our own findings.

The SO, characteristic of cortical LFP or EEG signals during NREMS, reflects the rhythmic alternation of active (UP) states, during which neurons are depolarized and show maximal firing, and silent (DOWN) states, periods of hyperpolarization and relative quiescence. The SO UP-state is thought to group the occurrence of δ waves and spindles in the cortico-thalamic network (Crunelli et al., 2018, Neske, 2015, Steriade et al., 1991). Our observation of δ2 waves nesting in almost all δ1 waves, thus adds further credibility to δ1 representing the SO. Furthermore, we show that δ2 is not only organized in time by δ1 waves but, at the level of EEG, contributes in turn to the multi-peak appearance of the latter. The two δ sub-populations preserved their characteristic waveforms throughout recovery, as did the bi-modal distribution of all SWs. Furthermore, the observation of a “slowing down” of the overall spectrum over time (Fig. 2D), was a direct result of the decrease in incidence of these faster waves δ2.

### Neuronal substrates of δ1 and δ2

The two SW types, which we identified based on their sleep-wake driven dynamics, are likely to originate in specific brain areas and generated through distinct mechanisms. To date, such dynamics have rarely been taken into account when studying the cellular and network substrates of SWs generated in the thalamocortical circuitry typical of NREMS. Directly relating our findings to *in vitro* work or studies using anaesthetized, awake, or immobilized animals recorded out of a dynamic sleep-wake history context might therefore not always be straightforward. We found that the well-known frontal dominance of EEG δ-power increases after SD (Cajochen et al., 1999, Munch et al., 2004), was specific to the δ2 band in both species, whereas δ1 increases were smaller and concerned a larger percentage of the cortex, a finding also observed recently in humans (Bersagliere et al., 2018). This latter observation is consistent with the SO engaging the entire neocortex (Contreras and Steriade, 1995, Massimini et al., 2004, Neske, 2015). Indeed, the SO was first described as a cortical phenomenon with layer 5 pyramidal neurons being key to UP-state initiation (Beltramo et al., 2013, Contreras and Steriade, 1995, Neske, 2015), while for δ-waves two sources have been described, one a clock-like oscillation generated by the intrinsic properties of thalamocortical neurons when hyperpolarized (McCormick and Pape, 1990, Steriade et al., 1991), and another of cortical origin likely to be generated in a similar fashion to cortical UP and DOWN states characteristic of the SO (Neske, 2015). It has become increasingly clear that thalamocortical crosstalk is an important contributor to the generation of both the SO and δ-waves (Crunelli et al., 2015, Neske, 2015). For instance, cortical UP-states synchronize thalamic δ-oscillations (Steriade et al., 1991), which might contribute to the prominent nesting we observed of δ2-waves at δ1 onset, while excitatory thalamocortical input to the cortex can trigger cortical UP-state initiation and removing thalamic input reduces cortical SO period and synchrony (David et al., 2013, Lemieux et al., 2014, Gent et al., 2018a). Moreover, thalamic neurons are intrinsically capable of generating rhythmic oscillations at <1Hz frequencies (Blethyn et al., 2006, Crunelli et al., 2018, Fernandez et al., 2018, Halasz et al., 2014, Herrera et al., 2016) that could aid the strengthening of the cortical SO. These and other observations demonstrate that SWs must be regarded as an emerging property of the thalamocortical network acting as a single unit (Crunelli et al., 2015). Consistent with this, our *in vivo* LFP recordings showed that the dynamics typical of each of the two SW populations could be observed throughout much of the cortex and thalamus with little variation precluding identification of their primary respective source.

In an effort to determine a thalamic contribution to either SW population, we optogenetically inhibited the centromedial thalamus (CMT), a non-sensory thalamic nucleus that we found to be important in the timing and manifestation of NREMS cortical SWs (Gent et al., 2018a). CMT inhibition boosted δ2 activity specifically, leaving δ1 in the cingulate cortex unaffected. Whether the δ2 augmentation is of thalamic or cortical origin is unclear as we quantified SWs during NREMS episodes consecutive to those during which the CMT was optogenetically silenced, indicating that increases in δ2 activity might be a cortical rebound phenomenon as SWs were suppressed during preceding optogenetic silencing (Gent et al., 2018a). Alternatively, optogenetic silencing of CMT neurons might have affected local circuitry dynamics recruiting other thalamic nuclei into modulating δ2 activity, or the suppression of SWs during CMT silencing could have released δ2 waves from the constraints imposed on their occurrence by δ1 waves (i.e., nesting). Optogenetic stimulation of inhibitory thalamic reticular neurons and pharmacological inhibition of the excitatory somatosensory thalamus (Lewis et al., 2015, Poulet et al., 2012), were both shown to induce cortical SWs at δ2 frequencies, confirming our CMT results, although it should be noted that these results were obtained in awake mice. Thus, our current data do not allow for the dissecting of precise elements in the thalamocortical circuitry underlying the δ1 and δ2 generation. Outside of the thalamic involvement we established, a cortical contribution, especially for δ1 activity, is more than likely. Our data also underline the need for studying SW generation in a dynamic and *in vivo* context, to better elucidate the cellular and network substrates of the sleep-wake driven changes in SWs.

### The paradox of coexisting deep NREMS with wake-like physiology

The high δ2 levels reached during NREMS immediately after SD were accompanied by high levels of a number of (neuro-) physiological variables more reminiscent of waking than of NREMS, suggesting that during this time it carries signatures of preceding wakefulness setting it apart from subsequent sleep episodes. Paradoxically, these early NREMS episodes in mammalian sleep are the deepest and considered most “recuperative” (Davis et al., 1937, Rechtschaffen et al., 1966).

The process of transitioning from wake to sleep is a phase characterized by a short-lived (several minutes) coexistence of wake- and sleep-like EEG patterns in the forebrain, indicating that the timing of ‘falling asleep’ varies among brain structures, with the thalamus becoming deactivated prior to the cortex, and fronto-central cortical areas prior to more posterior regions (Fernandez Guerrero and Achermann, 2019, Marzano et al., 2013, Nobili et al., 2012, Siclari et al., 2014, Siclari and Tononi, 2017). Our data extend these observations to physiological variables such as muscle tone, heart rate, and temperature during SD recovery and also demonstrate that under these conditions this transition phase is prolonged, encompassing the entire first NREMS episode in humans and the time up to the “pivot point” in mice. This is strikingly illustrated in the mouse where NREMS with the highest SW activity (i.e., “deepest sleep”) coexists with exceptionally high brain temperature, muscle tone, and beta/low-gamma EEG levels, all atypical for this state.

One contributing factor extending the transitional NREMS phase after SD is that high sleep pressure shortens sleep-onset latency thereby curtailing quiet waking preceding sleep onset, and during which, under non-SD conditions, much of the decreases in physiological variables can already occur. Numerous other processes coinciding with the wake-to-sleep transition could contribute to the aftereffects of extended waking. For example, although its precise dynamics still need to be determined, restoration of extracellular ionic balance in the brain, which can differ among brain areas and locally affect EEG SW activity (Ding et al., 2016), may take longer after extended periods of wakefulness.

We propose that these radical changes across a short period of time, reflect a transient NREMS state. The co-existence of incongruous sleep- and wake-like features during this phase is, in some respects, reminiscent of NREMS parasomnias, including sleep-walking (somnambulism), when wake behaviors most often occur during the first NREMS period when sleep is deepest and, at the level of the EEG, is associated with increases in δ activity in frontal brain areas and increases in wake-like EEG activity in the alpha and beta bands in sensorimotor areas (Castelnovo et al., 2018, Flamand et al., 2018, Nobili et al., 2011). Thus, the high initial levels of δ2 and subsequent steep decline we observed might not only be emblematic of this transient NREMS phase, but also instrumental in facilitating the passage from a highly active brain state and physiology to typical NREMS. Through ‘deactivating’ the thalamus and frontal cortex, this would prevent conscious experience of a still awake system, as is the case during sleep-walking. The finding that EEG δ-activity during anesthetic-induced loss-of-consciousness shares features with those of early NREMS after SD, is consistent with this line of thought (Baker et al., 2014, Zhang et al., 2015).

### Implications for models on sleep regulation and function

The dynamics of the sleep homeostatic process, aka *Process S* in the two-process model of sleep regulation (Borbely, 1982, Daan et al., 1984), were derived from the sleep-wake dependent changes in EEG δ power during undisturbed NREMS. This accessible and highly predictive EEG variable has not only been instrumental in understanding the regulation of sleep-wake timing but was also used as the basis for theories on sleep function, as it reflected ‘sleep need’ (Krueger et al., 2008, Tononi and Cirelli, 2006). Its acceptance is so wide-spread that EEG δ power has now become equivalent to the underlying sleep homeostatic process and any interventions affecting the manifestation of SWs are taken as proof of altered sleep homeostasis.

The dynamics of the full δ band show gradual and continuous changes in relation to sleep-wake distribution and can be reliably modelled by assuming exponential saturation functions for the increase during wake and decrease during NREMS. Here we show that this gradual time course is an artefact of combining the EEG activity of two SW populations with very different sleep-wake driven dynamics. Because δ1 activity was not in a quantitative relationship with SD duration, did not increase after spontaneous wakefulness, and did not decrease (in humans) or even increased (in mice) during initial recovery NREMS, it cannot be considered a proxy of an underlying sleep homeostatic process. Nevertheless, δ1 activity was increased after SD, as compared to baseline, and decreased over the remainder of sleep due to changes in δ1-wave incidence but not amplitude. However, this increase in δ1 after SD in the mouse could relate to aspects other than extended wakefulness, such as the intervention to keep mice awake. Furthermore, the δ1 level reached after SD in humans did not differ from baseline. Conversely, δ2 power and δ2-SW incidence and amplitude, all increased as a function of time-spent-awake and decreased during recovery sleep, suggesting a homeostatic mechanism. δ2 activity measures were, however, depleted within one hour, belying its functional necessity to continue sleeping.

Sleep-wake dependent changes in SW properties such as power and slope, are integral to the current leading hypothesis on sleep function stipulating that wakefulness is accompanied by synaptic strengthening that needs to be homeostatically balanced during NREMS through a process referred to as synaptic downscaling (Tononi and Cirelli, 2006). With the authors proposing SW-slope to reflect synaptic strength (Riedner et al., 2007, Vyazovskiy et al., 2007). We found that increased amplitude and incidence of δ2-waves fully accounted for the changes in SW-slope observed after SD with close to 1.0 correlations between slope and amplitude for SWs >2.0Hz (Fig. 2C). Similar findings were observed after SD in humans albeit for SWs >1.0Hz (Bersagliere and Achermann, 2010), consistent with the 1Hz lower δ1-δ2 demarcation in humans versus mice. If δ2 activity were to reflect a synaptic downscaling process, then it is accomplished within one hour. Although we cannot establish whether the process requires this little time and might indeed be an important function of this initial phase of sleep, the similarity of the discontinuous time course of δ2 with that of physiological variables not associated with the sleep homeostatic process makes it more plausible that these changes are part of transition phase before entering a more consistent resting state.

The discovery of two populations of SWs and their respective sleep-wake dependent dynamics concomitant to changes in a number of physiological variables, implies the existence of a transient NREMS state representing a mere ∼3% of total NREMS in the mouse across a 96-hour experiment. We believe that this early and fleeting portion of NREMS, which is enhanced by sleep deprivation, serves as a nexus between an active waking state and subsequent quiescent NREMS. The large, wake-dependent increases in δ2 activity during this transition phase could still serve a homeostatic function insomuch as it gates the transition into sleep. Such a scenario would allow us to refocus attention on the regulation and function of time-spent-in-NREMS and REM sleep, as their rebounds after SD have not been taken into consideration in the two-process model, nor in current hypotheses on sleep function, and which do not depend on δ power (Franken et al., 2001, Franken et al., 1991).

## Materials and Methods

### Mice experiments

#### Animals

All mice were C57BL/6J males, aged 10-16 weeks at time of implantation and were housed individually under 12-h light/12-h dark conditions. Food and water were given ad libitum. All animal procedures outlined were carried out using the guidelines of Swiss federal law and were preapproved by the Cantons of Vaud and Bern Veterinary Offices.

#### EEG/EMG/Thermistor implantation

Surgeries were performed according to previously published protocols under deep anesthesia using Ketamine/Xylazine solutions (Mang and Franken, 2012). In 6 mice (Figure 1A, B), electrodes were placed along an isolateral (right hemisphere) axis on the surface of the cortex (Bregma < 0.2mm) in the frontal (AP = +1.5mm, ML = +1.5mm), central (AP = −1.0mm, ML = +2.5mm), and parietal (occipital; AP = −3.0mm, ML = +1.5mm), brain regions (Franklin and Paxinos, 2013). All signals were differentially referenced with an additionally electrode implanted above the cerebellum. Subsequent experiments and datasets used only frontoparietal electrodes (n=32). EMG wires were implanted bilaterally in the cervicoauricularis muscles of the neck differentially recorded with one another. Prior to the experiment, animals were given 72-hours to recover from implantation, with a further 10 days to habituate to the recording cable. EEG/EMG signals were acquired using a commercially available system (Embla; Medcare Flaga, Thornton, CO, USA). Briefly, signals were amplified, filtered, and analog-to-digital converted to 2000Hz and downsampled to 200Hz for analysis. A separate group of mice (LFP, optogenetics experiments) were recorded using the Intan RHD2000 signal processor which sampled all channels at 20000 Hz. Signals were then downsampled to 200 Hz and analyzed in Matlab using custom scripts (see below), and scored in Somnologica (Medcare Flaga, Thornton, CO, USA) using identical criteria. In a subset of mice (n=9) a thermistor was inserted between the frontal and parietal electrodes to record cortical surface temperature (Hoekstra et al., 2019). A thermistor (series P20AAA102M, Thermometrics, Northridge, CA) was inserted through the skull on the surface of the right cortex (2.5 mm lateral to the midline, 2.5 mm posterior to bregma). Temperature was determined based on voltage levels recorded by the acquisition system, by measuring changes in the resistance of the individual thermistors based on manufacturer-supplied references values, were given a constant current of 100µA. For details on integrating constants and converting to °C, see (Hoekstra et al., 2019).

#### Discrimination of sleep wake states by EEG/EMG

Following acquisition, signals were filtered for power-line artifacts resulting from AC power cycling (50 Hz). Scoring of sleep and wake states in 4 second epochs was achieved using previously published criteria (Mang and Franken, 2012), and classified as either waking, non-rapid eye movement (NREM) sleep, or REM sleep, using visual inspection without knowledge of the recording condition.

#### Thermoelectric generator (TEG) installation

To control levels of brain temperature during SD recovery, in a subset of mice (n=4) an aluminum plate (taken from a commercially available CPU heatsink: Intel BXSTS200C) was placed on the skull of the animal contralateral to the thermistor (see above, and Figure 5G), using thermal paste (WLP-1, S+S Regeltechnik GmbH) to ensure efficient temperature transfer. A thermoelectric generator (Peltier module: Laird Thermal Systems, Inc. 45850-503) was then affixed using dental cement (Paladur) 2 cm above the animal and affixed to the EEG/EMG recording cable. Temperature manipulation was achieved using a Bench Top Power Supply (RND 320-KD3005P) to control amperage input to the TEG. Output temperatures were calibrated prior to the experimentation. Thermal images were acquired using an infrared thermal camera (Model E30, FLIR Systems), and analyzed using FLIR tools software (FLIR Systems).

#### LFP tetrode implantation

Tetrodes were constructed in lab using 4 strands of 10µm twisted tungsten wire, which were then attached by gold pins to an electrode interface board. They were then placed in 5 thalamic and cortical structures: CMT (central midline thalamus) (AP −1.7 mm, ML +1.0 mm, DV −3.8 mm, 15°), CING (cingulate cortex) (AP +1.8 mm, ML +0.2 mm, DV −1.6 mm), AD (anterodorsal thalamic nuclei) (AP −0.9 mm, ML ± 0.8 mm, DV −3.2 mm), BARR (barrel cortex) (AP −2.0 mm, ML +2.2 mm, DV −1.1 mm) and V2 (visual cortex) (AP −3.3 mm, ML +2.5 mm, DV −0.9 mm) and secured to the skull with dental acrylic (C&B Metabond). Optic fibers of 200µm diameter were placed in the CMT (AP −1.7 mm, ML +1.0 mm, DV −3.8mm, 15and secured with the same dental acrylic. Finally, the implant was stabilized using a methyl methacrylate cement and the animal allowed to recover in the home cage on top of a heating mat. Animals were allowed a minimum of 5 days to recover before starting recordings.

#### Optogenetics experiments

C57Bl6 mice, aged 6 weeks were anaesthetized using isoflurane (1.0–1.5% with oxygen) and placed in a stereotaxic apparatus (Model 940, David Kopf Instruments). Injections of AAV were performed using a 10-µL Hamilton syringe attached to an infusion pump (Model 1200, Harvard Apparatus), in the CMT (AP –1.7 mm, ML +1.0 mm, DV –3.8 mm, 15°, 100 nL), based on coordinates from the mouse brain atlas (Franklin and Paxinos, 2013) and performed at 0.1 µL/min and the needle left in situ for 10 min afterwards to facilitate diffusion. Animals were injected with AAV2-CaMKII-E1fa-ArchT3.0-EYFP for optical silencing of CMT neurons. All plasmids were obtained from University of North Carolina Vector Core Facility. Animals were given 21 days to recover before optogenetic experiments commenced. For more details see (Gent et al., 2018a). Animals were instrumented for tetrode recording (see above) 3-4 weeks after virus injection to allow sufficient time for opsin expression.

Optical fibers connected to a black furcation tubing coated patch chord (Doric Lenses) were further covered in black varnish to reduce optical leakage from the laser. Animals were habituated to being tethered for up to 8 h per day until a normal sleep–wake episode resumed based on EEG recordings from ZT4-9. Experiments were performed from ZT4-8, following SD. Inhibition was achieved with a green laser (532 nm; LRS-0532-GFM-00100-03, Laserglow Technologies starting 10 s NREMS onset. Output was controlled using a pulse generator TTL (Master-9, AMPI or PulsePal 2, Sanworks), co-acquired with all recordings. During sleep deprivation, animals were stimulated with either blue or green lasers (for stimulation/inhibition), during specific NREMS episodes. The episode immediately following was then analyzed for spectral differences. For more details see (Gent et al., 2018a).

#### Sleep deprivation protocol

After >10d of recovery from surgery mice were recorded for EEG/EMG and LMA for a total of 96 hours. For the first 48h animals remained in undisturbed baseline conditions. Beginning on the third day, mice underwent a 6h sleep deprivation starting at light onset (ZT0) using gentle handling (Mang and Franken, 2012). Animals were then allowed to recover for a further 42 hours. Other experiments presented in this article also involved SD starting at different ZT across the 24-hour period (Fig. 3C, (Curie et al., 2013)), or of varying lengths (2 and 4h, Fig. 3B).

For LFP recordings and optogenetic experiments, animals were taken from their home cages and placed in new ones at ZT0 with clean bedding, food, and water, in addition to a novel plastic object. A 4-hour sleep deprivation was performed before mice were transferred to their original cages for data acquisition between ZT 4 and 9. For more details see (Gent et al., 2018a).

#### Respiratory recordings

To determine the frequency of breathing, mice were simultaneously recorded for EEG/EMG in conjunction with a piezoelectric system (Signal Solutions, LLC, Lexington, KY, USA) which uses breathing-related movements to estimate sleep-wake state (Mang et al., 2014). Briefly, the piezoelectric platform comprises a polycarbonate cage and a floor covered with a polyvinylidine difluoride (PVDF) film (17.8 × 17.8 cm, 110 μm thick (Measurement Specialties, Inc., Hampton, NY) covered with standard rodent litter. A 6-feature vector is extracted from the piezoelectric signal for each 4-seconds, which was matched to EEG/EMG derived sleep and wake states. Respiratory rate is one of the vector features. For further details see (Yaghouby et al., 2016).

#### Frequency-domain analysis of EEG sleep/wake signals

Power spectral density for each 4-s EEG window were generated using a discrete Fourier transform after using a Hamming window, yielding power density spectra (0 to 100 Hz) with a frequency resolution of 0.25Hz. Bins containing frequencies between 49-51Hz were excluded due to intrinsic power-line artifacts in some animals. Epochs containing signal artifacts were separately identified to be included in state quantification though not in spectral analysis. Of note, frequencies are annotated based on midpoint of 0.25 Hz bin (e.g. 2Hz = 1.875 to 2.125 Hz). δ power was calculated for 4-seconds of NREMS surrounded by artifact-free epochs of the same behavioral state. Consolidated NREMS episodes were defined as uninterrupted sequences of at least 32s (8 epochs), based on visual inspection and previous publications (Mang and Franken, 2012, Vassalli and Franken, 2017).

#### Time-Course Analysis of spectral power in NREMS

The spectral band (0.25-90Hz), was separated into its components to examine dynamics across time of individual bands: δ (1-4Hz), δ1 (0.75-1.75Hz), δ2 (2.5-3.5Hz), theta (6-9 Hz), sigma (10-15 Hz), and low gamma (32-47 Hz). To control for interindividual differences, the absolute spectral power of the each of these frequency bands, was expressed to values calculated to the period during the two baseline (12hL:12hD) days with the lowest mean value, corresponding to the last 4h of the light period (ZT8–12), according to previously published methods (Franken et al., 1999, Mang and Franken, 2012). Values were averaged into time periods of equal length for baseline light (12 percentiles), dark (6 percentiles), and following SD [either 8 (Fig. 1) or 25 percentiles (**others**)].

#### Time-course analysis of δ1 and δ2 dynamics across Wake/REMS-to-NREMS transitions

Following SD, the dynamics of the sub-bands (δ1 and δ2), were analyzed per epoch across transitions from either wake or REMS, to NREMS, similar to previously published methods (Franken et al., 1998, Vassalli and Franken, 2017). Transitions were separated into 4 groups, the first 3 percentiles calculated starting at SD recovery, and the ensuing hours following the pivot point (see section below). A Wake/REMS-to-NREMS transition was defined as ≥4 consecutive 4s epochs of NREMS, preceded by ≥8 scored as either wake or REMS. δ1 and δ2 spectral power was calculated across these transitions starting at 1mn before and 2mn after, and then averaged across these subsequent 3-minute windows. Values were expressed relative to changes during two baseline recording days (ZT8-12), as for δ power. Rise-to-maximum curves for the first percentile were used to illustrate the rapid ascent of δ2 using SigmaPlot (v. 12.5 SysStat 2011).

#### Slow-wave slope analysis

Raw EEG signals at 200Hz were imported into Matlab (v. 2018, Mathworks Inc.), in addition to sleep/wake scores. EEG were then filtered for δ spectrum (Chebyshev type-II 0.5-4.5 Hz, bandstop at 0.1 and 10 Hz), similar to others (Freyburger et al., 2017, Panagiotou et al., 2017, Vyazovskiy et al., 2007, Massart et al., 2014). To detect slow-waves, a custom-made Matlab algorithm was employed, based on zero-crossings and wave reconstruction closely following others (Freyburger et al., 2017, Riedner et al., 2007). Detection of NREMS SWs was achieved using the following steps: (1) zero-crossings were determined, (2) surrounding local maxima and minima were identified, (3) thresholding was applied to control for SW amplitude (see below), (4) mathematical slope based on amplitude and SW frequency (period) was determined in all possible directions and combinations (see Fig. S1). The slopes which were analyzed in the manuscript refer to zero-crossing to maximum amplitude SWs which begin with a negative deflection. Though all slopes were calculated (Fig. S1), their mean relative changes across time were nearly identical. Data was then grouped and averaged for NREMS episodes longer than 32 seconds. Slope, amplitude, and period changes were expressed as a percentage of baseline ZT8-12 NREMS values, as with spectral power.

The above algorithm detected SWs across the entirety of the 96-hour experiment for all NREMS episodes lower and upper thresholding was applied to capture “true” NREMS SWs based on visual inspection and previously published criteria (Freyburger et al., 2017, Vyazovskiy et al., 2007). Upper thresholds were set at 6 times the SD of the amplitude of all detected waveforms. The lower threshold was set at the 95% probability limit of SWs detected during REMS, with the idea that SWs would not appear during REMS. For amplitude distribution, see Fig. S1D. Following SW detection, SWs were categorized according to their frequencies in 0.25Hz windows and analyzed for incidence and amplitude (see Fig. 2 A, B & Fig. S3 A, B).

Average waveforms during NREMS episodes (Fig. S3 C, D) were first located using the filtered EEG signal for δ1 and δ2, and then interpolated to fit 800ms (δ1) and 300ms (δ2), to better compare individual SWs of different frequencies within these ranges. The timecodes of these SWs were then reutilized to extract raw unfiltered waveforms. To determine onset and offset of nested δ2 waves, locations of δ1 waveforms were stored and the raw signal was then filtered only for δ2 (2.5-3.5Hz) using the same Chebyshev filter characteristics (see above). δ2 waveforms which fell within these timecodes were considered “nested” and their onset/offset from δ1 waveforms was calculated.

### Analysis of physiological variables

All variables presented in Fig. 6 and S5 (with the exception of heart rate), were expressed in relative values and grouped into identical percentiles as δ1 and δ2 (Fig. 5A). In mice this was 25 percentiles (5 prior to pivot point, and 20 after). Specific analyses are as follows:

- **Higher frequencies** (Fig. 6B) are calculated identically to δ power, except at different frequencies (18-45 Hz).
- **Muscle tone** (Fig. 6C) is calculated as the relative changes in mean EMG variance across NREMS episodes. EMG variance (σ2) is calculated based on the sum of the squared distances of from the mean, divided by the number samples (800). Values are expressed relative to EMG variance during NREMS of the baseline (ZT8-12).
- **Brain temperature** (Fig. 6D) is represented as differences during NREMS percentiles from average values during the 6-hour waking period (SD) preceding the recovery.
- **Respiration** (Fig. 6E) is described in a previous section and is expressed in absolute frequency.

### Hierarchical clustering to determine δ band separation

Cluster analysis of referenced (baseline ZT8-12) power per 0.25Hz, as in Fig. 2D, was achieved based on mean values from 38 mice for the first 8 percentiles of NREMS during SD recovery (see main text). The Matlab function “clustergram” was used to generate a dendrogram and heat map, using hierarchical clustering based on Euclidean distance metric and average linkage.

### Detection of UP/DOWN states and neuronal firing rates

Detection of UP/DOWN states was achieved using bandpass-filtered (0.5–3 Hz) LFP/EEG signals, in forward and reverse directions (filtfilt, Matlab). Individual UP/DOWN states were detected using zero-crossing method of these signals. UP state onset was defined as crossing from negative to positive, and those UP/DOWN states with an amplitude < 1 s.d. from the means, and shorter than 200 ms, were excluded. To extract multiunit activity from the LFP, signals were bandpass-filtered (600-4,000-Hz, fourth-order elliptic filter, 0.1-dB passband ripple, –40 dB stopband attenuation), and detected using a threshold of 7.5x the median of the filtered signal’s absolute value. Single units were then identified after sorting with the WaveClus toolbox (Quiroga et al., 2004). For more details see (Gent et al., 2018a). Average firing rate was calculated based on the number of spikes present per second of NREMS episode and expressed in Hz.

### Human experiments

#### Study participants

A total of 110 healthy young men participated in the studies used for this analysis. No participant had traveled more than two time zones in the preceding 3 months, nor suffered from a diagnosed nervous system disorder or other acute medical condition. Additionally, subjects were pre-screened in the laboratory prior to the study to confirm the absence of any sleep disorders and were free of medication and recreational drug use. All subjects signed informed consent and were compensated financially for participating. For more details see previous publications (Bodenmann et al., 2009, Holst et al., 2017, Valomon et al., 2018, Weigend et al., 2019). Some subjects were removed across 5 studies for signal artifacts, which made time-course analysis of spectral bands difficult. For topographical and RR interval analyses, one subject was removed for signal inconsistencies.

#### Data acquisition

Continuous recording of EEG, EOG, EMG, and ECG data was acquired during baseline and following a sleep deprivation. Sleep and waking stages were visually scored in 20-s epochs (C3-A2 derivation) according to standard criteria (Berry et al., 2012), using Rembrandt® Analysis Manager (version 8; Embla Systems, Broomfield, CO, USA). Movement- and arousal-related artifacts were visually identified and eliminated from subsequent analysis. Analog signals from each derivation (see Figure 5C), sampled at 256 Hz were filtered [high-pass (−3 dB at 0.15–0.16 Hz) and low-pass filtering (−3 dB at 67.2 Hz). EEG spectra were calculated identically as in mice (see above) using Matlab for each 4s epoch and averaged according to the assigned sleep-wake state, for each derivation available. NREM/REM sleep cycles were first defined according to (Feinberg and Floyd, 1979), and for BL and REC conditions, δ power time-course during all scored epochs was visually inspected to separate the first two episodes which did not have a long REMS episode between them. Only the first 8h of recovery sleep was analyzed. EEG power was expressed to all-night averages of the baseline period. Each NREMS episode was separated into 10 equal percentiles. To compare relative changes across the scalp in a subset of individuals (Fig. 5C, D), mean power across all sites for each δ sub-band during baseline was used. Mean power for electrode site during SD recovery was then estimated at the maximum percentile during for δ1 and δ2 during each NREMS episode and expressed as a log2 ratio to baseline.

#### Statistical analyses

All statistical analyses were performed in either Statistica 8.0 (Statsoft, inc) or using built-in MATLAB functions. Sleep-wake distribution, EEG spectral power, and time-course dynamics of specific frequency bands were assessed using 2- or 3-way repeated-measures analysis of variance (rANOVA). Statistical significance was considered as p < 0.05, and all results are given as mean values ± SEM. Tukey’s post hoc test was used to determine significant effects and interactions and corrected for multiple comparison. Comparison of two-groups was achieved with Student’s t-test. Statistical methodology is further described in the results section and figure legends. Linear and piece-wise regressions were performed, and squared Pearson’s correlation coefficients were statistically compared through Fisher Z transformation followed by t-test.

#### Data and code availability

The code supporting the current study have not been deposited in a public repository because of their specificity to file structures but are available from the corresponding author on request.

## Supplementary Figures

**Figure S1:**
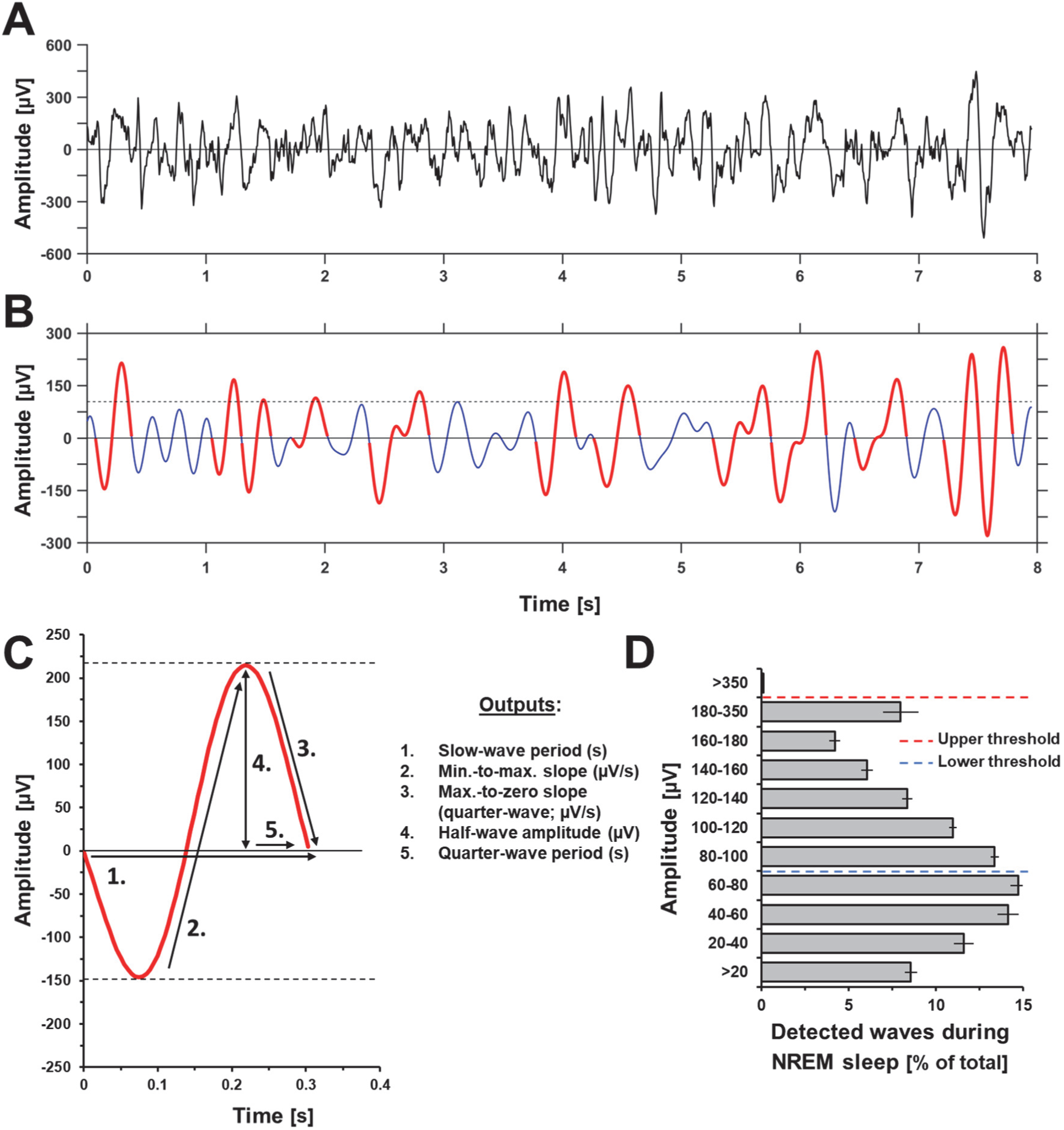
SW-slope detection algorithm. (**A**) 8-second raw EEG trace during NREMS. (**B**) EEG signal was filtered using a Chebyshev Type II filter (**see Methods**), and a zero-crossing algorithm was employed to detect slow waves (SWs)s. Waves of higher amplitude than threshold [individual mean maximum amplitude (95%) of waveforms during REM sleep; dashed line; lower threshold in D] were considered SWs (highlighted in red). A maximum threshold was set at 6 times the standard deviation of the positive amplitude of the filtered signal to remove signal artifacts. (**C**) Selected SWs were then analyzed for amplitude, period, and slope. (**D**) Distribution of waveforms by amplitude during NREMS expressed as a total of all detected waves across 48h of baseline and 42h of SD recovery, with maximum (red dashed line) and minimum (blue dashed line) threshold values (n=38). Determination of maximum and minimum thresholds is detailed in the Methods section.

**Figure S2:**
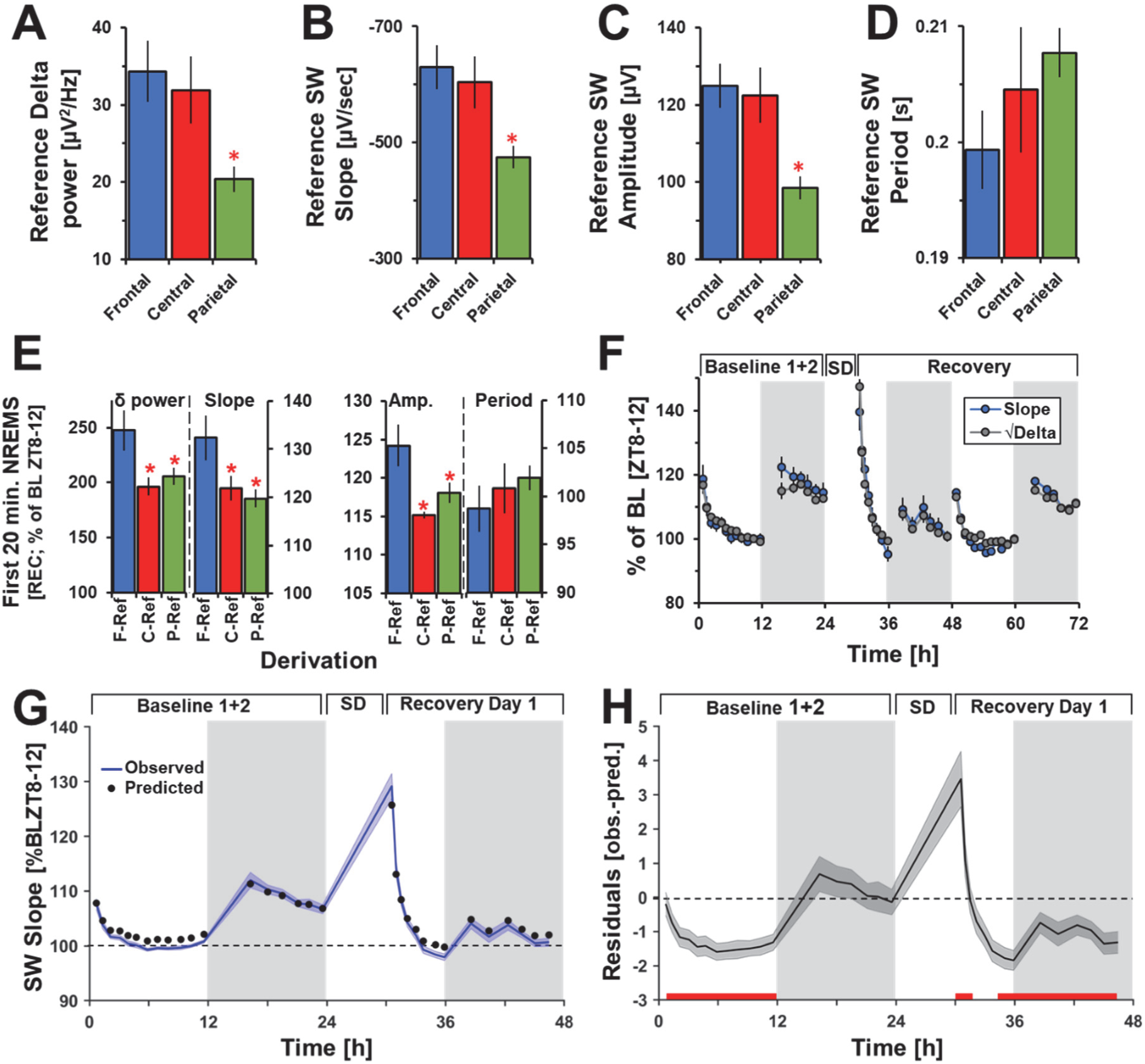
Time- and frequency-domain analyses illustrate site-specific and SD recovery-dependent difference in SW properties. Mean levels for the F-, C-, and P-Ref derivations during baseline (ZT8-12) of **(A)** δ power, **(B)**, SW-slope **(C)**, SW-amplitude, and **(D)** SW-period. δ power, SW-amplitude, and SW-slope are significantly affected by derivation, while SW-period is not (one-way ANOVA derivation: p_δ_ =0.01; p_slope_=0.007; p_amplitude_=0.01; p_period_=0.33). These values serve as the 100% values for all calculated dynamic changes (% of BL ZT8-12). (**E**) Initial relative changes (first 20 minutes) of SW parameters following SD. As in **A-D**, period is not significantly affected, with the largest values seen in the frontal derivation (one-way ANOVA derivation: p_δ_=0.02; p_slope_=0.03; p_amplitude_=0.008; p_period_=0.58, respectively). (**F**) SW-slope and the square-root of δ power for the F-P derivation across baseline and SD recovery were highly correlated (R^2^=0.93, p<0.00001, n=68). (**G**) Observed (blue line) vs. predicted (black circles) values of SW slope based on linear regressions with SW amplitude for each time percentile (R^2^=0.85, p<0.00001, n=32). (**H**) Residuals of the linear model (observed-predicted, see G), might relate to changes in SW period, the second component of SW slope. Values in **A-H** represent mean values +/- S.E.M (**A-E**: n=6; **F-H**: n=38) referenced to the baseline ZT8-12. Red asterisks (**A-E**) denote post-hoc significance with frontal derivation (p<0.05). Red bars (**H**) show significant deviations from zero (p<0.05). Values in B-D are given for quarter-waves from minimum to proceeding positive zero-crossing (see Fig. S1C).

**Figure S3:**
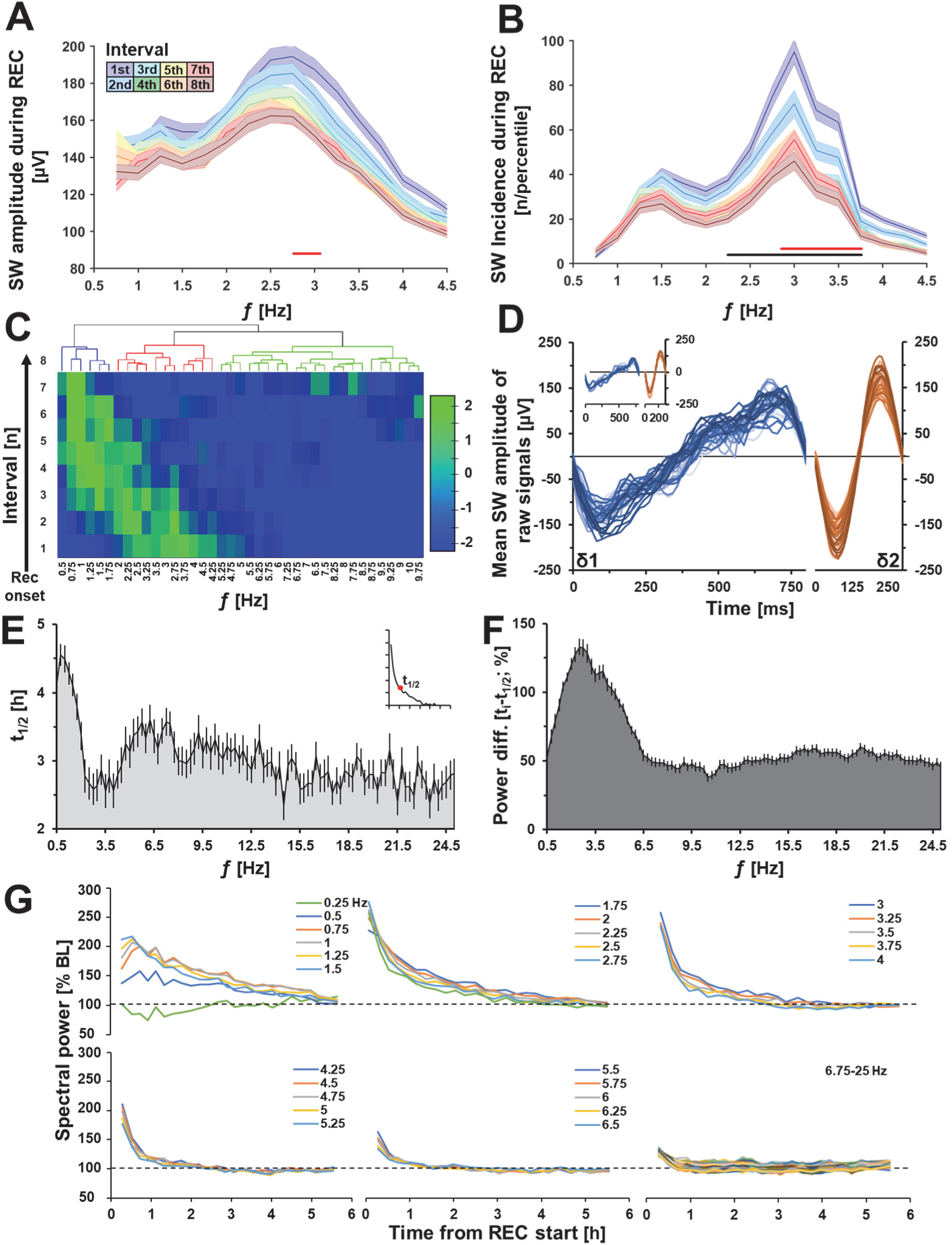
Defining the δ1 and δ2 frequency bands. (**A**) Mean SW-amplitude of filtered signals per 0.25Hz bin of detected SWs during the first 8 percentiles of SD recovery (as in Fig. 2A). (**B**) Uncorrected mean SW-incidence binned per 0.25Hz (as in Fig. 2B). (**C**) Unsupervised hierarchical clustering of the first 8 percentiles (as in **E**), grouping δ1 (0.75-1.75 Hz; blue) and δ2 (2.5-3.5 Hz; red) bands, and higher frequencies (4.75-10 Hz; green). Brighter green indicates closer association. (**D**) As in Figure 2F, average waveforms of unfiltered (raw) detected δ1 (**blue-left**) and δ2 (**orange-right**) waves during the first 10 minutes of recovery NREMS following SD, for individual mice in a frontal-parietal derivation (n=38). Note that the fronto-cerebellar derivation yields similar average waveforms (**insert**; n=6). (**E**) Time required for power density in the respective frequency bin to reach half of its initial value during SD recovery (t_1/2_). Higher frequencies (>4.5Hz) reach t_1/2_ faster than the slowest (<2Hz), however initial values start much lower and thus decay less overall (ratio difference; Fig. S3F) (**G**) Spectral power density for each 0.25Hz bin across the first 6 hours of recovery expressed in 25 approximately equal time percentiles. Higher initial values are observed as frequency increases within the δ band. Bars in **E** and shading in **A-B** represents s.e.m. (n=38); significant changes (rANOVA percentile x frequency p<0.001; post-hoc t-test p<0.05) between 1st-2nd and 2nd-3rd percentiles are indicated respectively in red and black bars below.

**Figure S4:**
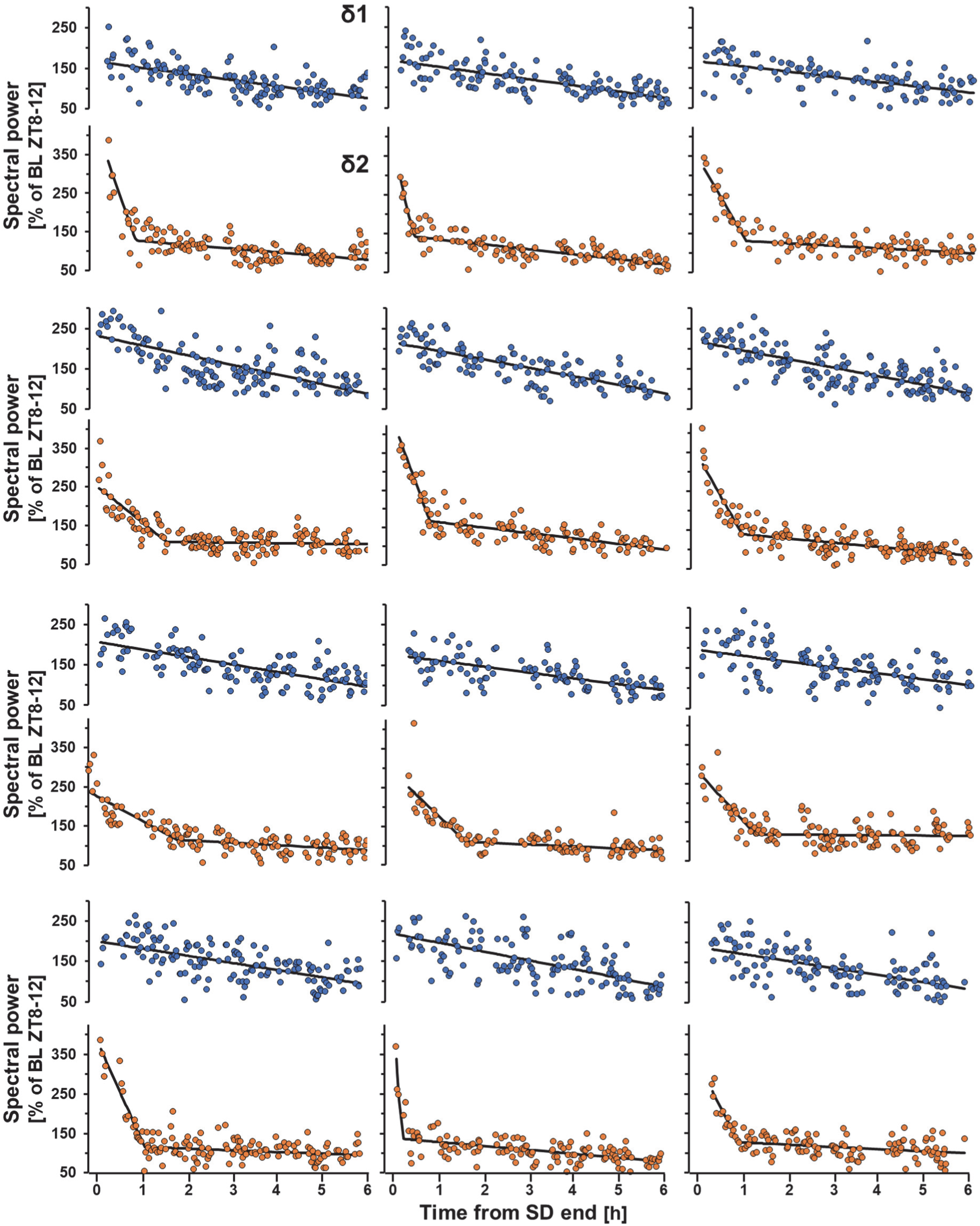
Individual NREMS episode plots for δ1/2. Examples of δ1 (**blue**) and δ2 (**orange**) means per NREMS episode (>32s) during the first 6 hours of recovery for 12 individual mice. Note the rapid decrease in δ2 for most mice during initial episodes, and the overall linear trend of δ1.

**Figure S5:**
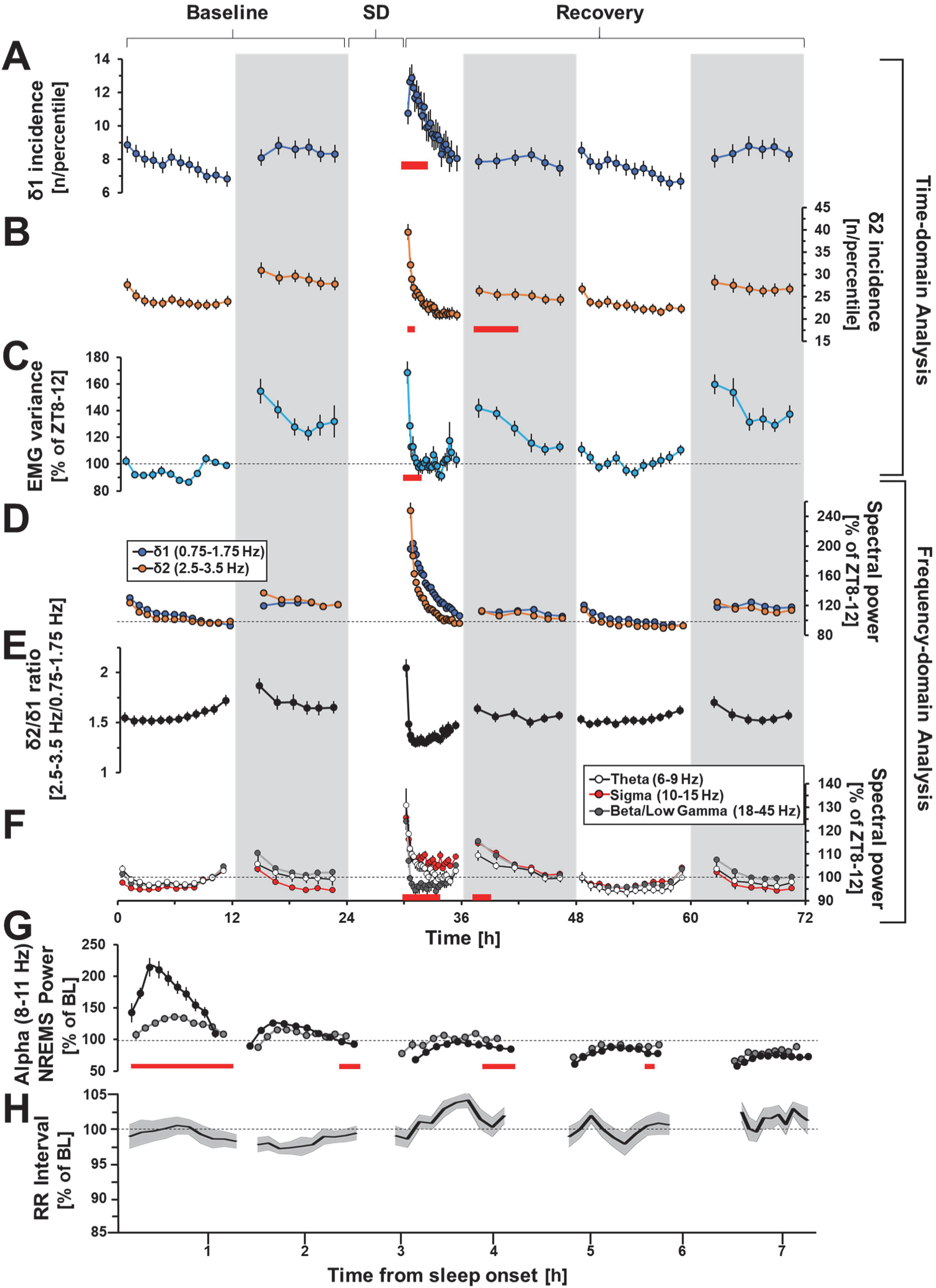
Initial changes across different measures consecutive to prolonged waking reflect a NREMS sub-state. Density of detected NREMS SWs within the δ1 (**A**) and δ2 (**B**) range across baseline and recovery. Note the similarity with the time dynamics of spectral power in the two bands (Fig. 6A). Of note δ2 but not δ1 density was lower in the first dark period of the recovery when compared to their corresponding baseline values (rANOVA time: p<0.0001). Increases in EMG variance (**C**), δ2/δ1 relative power (**D**), δ2/δ1 ratio (**E**), and other spectral components (**F**), increases are also seen during the baseline dark period after prolonged spontaneous waking (baseline, 2^nd^ recovery day), as in Fig 4A, 6A, during recovery. Initial values of recovery are significantly different from baseline for these three measures, as with δ2 density. (**G**) Alpha activity (8-11 Hz) during baseline (black circles) and recovery (grey circles) NREMS in humans. (**H**) RR interval during baseline NREMS in humans (see Fig. 6H). Red squares indicate significant post-hoc differences between baseline and recovery. Data is presented as mean values ± SEM.

**Figure S6:**
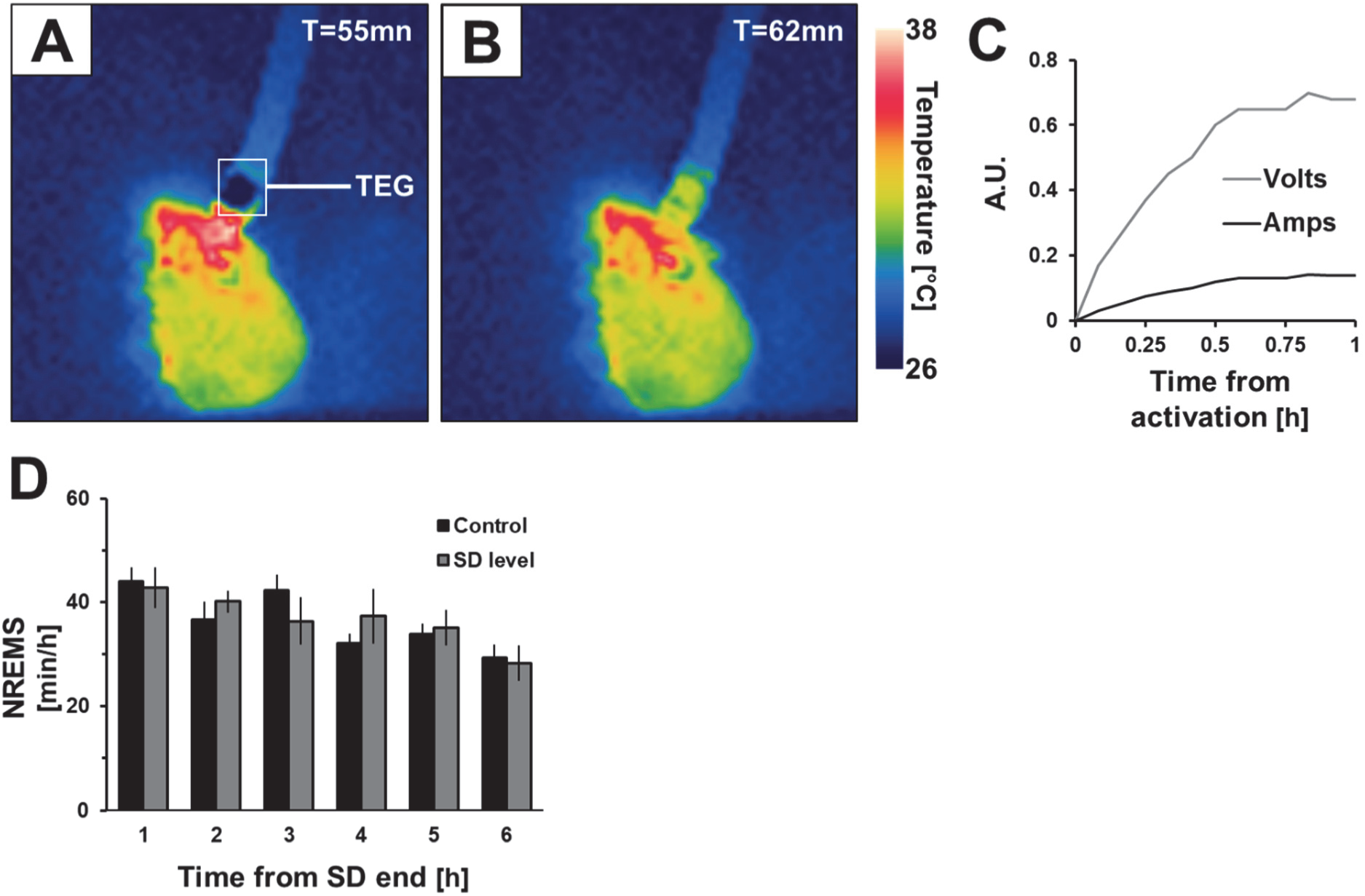
Confirmation of TEG efficacy during cortical warming. Thermal image of one mouse undergoing cortical warming during (**A**) and 2 minutes after the TEG has been deactivated (**B**). The cooled side of the TEG is visible above the EEG connector, which begins to reach ambient temperatures after deactivation. (**C**) Example in one mouse of the increase in power needed to compensate for decreases in brain temperature during the manipulation. (**D**) NREMS amounts for control and SD level conditions for the first 6 hours after SD. As stated previously, warming is only performed during the first hour of recovery.

## Acknowledgements

We would like to thank Diego M. Bauer for his assistance in preparing the human EEG data used in this study. Furthermore, we greatly appreciate the insights on this project and manuscript provided by Pr. Peter Achermann, Pr. Anita Luhti, and Dr. Laura Fernandez. This study was performed at the Universities of Lausanne, Bern, and Zurich, Switzerland, and supported by the Swiss National Science Foundation (SNF n°146694, Sinergia 136201) and the States of Vaud (supporting JH, MMBH, and PF), Bern (supporting TG and AA), and Zurich (supporting H-PL). Further support for JH was provided by the Dr. Rub Foundation, and the University of Lausanne Foundation. Current address for T. Gent: Vetsuisse Faculty, University of Zürich, Zurich, Switzerland. Current address for M.M.B. Hoekstra: Department of Brain Sciences, University UK Dementia Research Institute at Imperial College London, London, United Kingdom.

## Author Contributions

The study was designed by J.H., P.F., T.G., A.A., and H-P.L. P.F.; Experiments were performed by J.H., M.M.B.H, V.M., T.G., and Y.E.; J.H. and T.G. analyzed the data; J.H., P.F., and A.A. wrote the manuscript.

## Declaration of Interests

The authors declare no competing interests.

